# A novel subset of hepatocytes is simultaneously gluconeogenic and *de novo* lipogenic in the fed state and is naturally insulin resistant

**DOI:** 10.64898/2026.03.09.710526

**Authors:** Junichi Okada, Austin Landgraf, Maxwell Horton, Yunping Qiu, Alus M. Xiaoli, Roberto Ribas, Li Liu, Sofia V. Krylova, Hernando J. Olivos, Victor L. Schuster, Fajun Yang, Takeshi Saito, Ramon C. Sun, Meredith Hawkins, Gary J Schwartz, Carolina Eliscovich, Kosaku Shinoda, Irwin J. Kurland, Jeffrey E. Pessin

**Author notes:** Please address correspondences to: Junichi Okada, Department of Medicine (Division of Endocrinology), The Albert Einstein College of Medicine, 369 Price Center, 1301 Morris Park Avenue, Bronx, NY 10461.

## Abstract

It is generally accepted that hepatic gluconeogenesis, the synthesis of glucose from non-carbohydrate substrates is active in the fasted state and inactive in the fed state. In contrast, *de novo* lipogenesis is active in the fed state and is inactive in the fasted state. Here, we used targeted single cell RNA-seq, HCR RNA-FISH, and PrimeFlow in normal physiological mouse liver, and identified a subpopulation of periportal hepatocytes that simultaneously co-express both gluconeogenic and lipogenic genes in the fed state. Euglycemic-hyperinsulinemic clamps further demonstrated that this novel hepatocyte subpopulation is naturally insulin resistant. Spatial metabolic imaging coupled with stable isotope tracing analyses revealed individual hepatocytes that simultaneously undergo both gluconeogenesis and *de novo* lipogenesis. These dual-positive hepatocytes were also present in human hepatocytes from humanized mouse livers. Moreover, the number of dual-positive hepatocytes increased in high-fat diet-fed mice, suggesting a paradigm shift in our understanding of how the liver becomes insulin resistant.

## INTRODUCTION

The mammalian liver is composed of parenchymal (hepatocytes) and non-parenchymal (endothelial, Kupffer, cholangiocytes, stellate) cells organized into repetitive hexagonal arrays termed lobules^1,2^. Within each lobule the hepatocytes are distributed from the portal triad (portal vein, hepatic artery, bile duct) at the six vertices and the central vein located at the centroid of the hexagon. The hepatocytes aligned within a lobule display functionally distinct metabolic properties and are generally divided into three zones: the periportal hepatocytes (zone 1), midlobular hepatocytes (zone 2) and the pericentral hepatocytes (zone 3), with emerging single-cell studies indicating that hepatic zonation exists along a more continuous and finely partitioned spectrum^3,4^. The pericentral hepatocytes display higher rates of glycolysis, bile acid and glutamine biosynthesis^1,2^. The periportal hepatocytes are primarily responsible for beta-oxidation and urea oxidation and in particular, gluconeogenesis^1,2^. However, gluconeogenesis is a dynamic process that is regulated by numerous hormonal, neural and metabolic inputs. During early states of fasting *in vivo*, we found that periportal hepatocytes are initially responsible for gluconeogenesis, but as fasting duration increased, the mid-zone and then pericentral hepatocytes underwent increased gluconeogenic gene expression, protein levels and ultimately gluconeogenic activity that approached that of the periportal hepatocytes^5^. In contrast, lipogenic gene expression that is high in the fed state is rapidly downregulated to essentially undetectable levels during the initial time frame of fasting induction. These data demonstrate the remarkable plasticity of hepatocyte gluconeogenic gene expression and gluconeogenic activity across the liver lobule depending upon organismal nutrient state.

While it is generally accepted that *de novo* lipogenesis is activated in the fed state whereas gluconeogenesis and hepatic glucose output are activated in the fasted state, several studies observed a measurable rate of gluconeogenesis and hepatic glucose production in the fed state, despite the availability of dietary glucose^6,7^. Although not thoroughly investigated, the molecular and cellular bases for the basal rate of hepatic gluconeogenesis and hepatic glucose output was generally thought to represent glucose production for the indirect pathway of glycogen synthesis^8–11^.

In this study, we used a variety of single cell technologies to confirm the presence of basal liver gluconeogenic activity transcriptionally and functionally in the fed state that results in hepatic glucose release. Importantly, our data revealed that this basal gluconeogenesis results from a subset of unique hepatocytes that simultaneously display expression of both *de novo* lipogenic and gluconeogenic genes exclusively localized to the periportal region. Spatial metabolomic imaging further confirmed that this subset of periportal hepatocytes is capable of simultaneously undergoing both gluconeogenesis and *de novo* lipogenesis, and have high rates of ATP turnover, consistent with the energetic need to sustain these dual processes. Furthermore, euglycemic-hyperinsulinemic clamps and PrimeFlow analyses suggested that these hepatocytes are resistant to insulin suppression of gluconeogenic gene expression, demonstrating natural insulin resistance under normal physiological conditions. Moreover, the number of these naturally insulin-resistant hepatocytes is increased by high-fat diet feeding.

## RESULTS

### Gluconeogenic genes are transcribed in hepatocytes in the fed state

To investigate the regulation of liver gene expression in the normal physiological transitions between fed and fasted states, we used a previously established protocol in which mice rapidly consume food and fill their stomachs to create a relatively uniform fed state, and following food removal, facilitate a uniform transition into the fasted state^5,12^. We first confirmed our previous findings that lipogenic gene *Fasn* mRNA is highly expressed in the fed state of total primary isolated hepatocytes and nearly completely suppressed in the fasted state (Fig. 1A). In contrast to lipogenic gene mRNA, the gluconeogenic gene *Pck1,* was highly expressed in the fasted state, yet surprisingly there was a low but readily detectable mRNA in the fed state as well (Fig. 1B). There were no significant effects of fasting/feeding on the expression of constitutively expressed gene albumin (*Alb*) mRNA (Fig. 1C). To determine if the observed feeding state changes in mRNA levels were due to a transcriptional process, we examined the transcriptional activation of lipogenic and gluconeogenic genes by performing DNA-dependent RNA polymerase II (RNAPII) ChIP-seq to assess RNAPII gene occupancy in total primary isolated hepatocytes. As expected, RNAPII occupancy was relatively high across the gene body of *Fasn* in the fed state with very low binding in the fasted state (Fig. 1D). RNAPII occupancy on other lipogenic genes *Acly* and *Elovl6*, displayed a similar pattern (Fig. S1A and S1B). Both *Fasn* and *Acly* displayed a peak of RNAPII at the transcription start site indicative of proximal promoter pausing^13^. In contrast, the gluconeogenic genes *Pck1* and *G6pc* showed the opposite trend with higher RNAPII occupancy in the fasted state. Although both *Pck1* (Fig. 1E) and *G6pc* (Fig. S1C) displayed strong gene occupancy by RNAPII in their active (fasted) state, we also observed RNAPII gene occupancy in the inactive fed state. In addition, there was no evidence of proximal promoter pausing for the gluconeogenic genes. In contrast, there were no changes in RNAPII occupancy for *Alb* (a constitutive hepatocyte gene, Fig. 1F). As a control, the *Ins1* gene, which is not expressed in hepatocytes, showed virtually no RNAPII occupancy in either the fed or fasted state (Fig. 1G).

**Figure 1.**
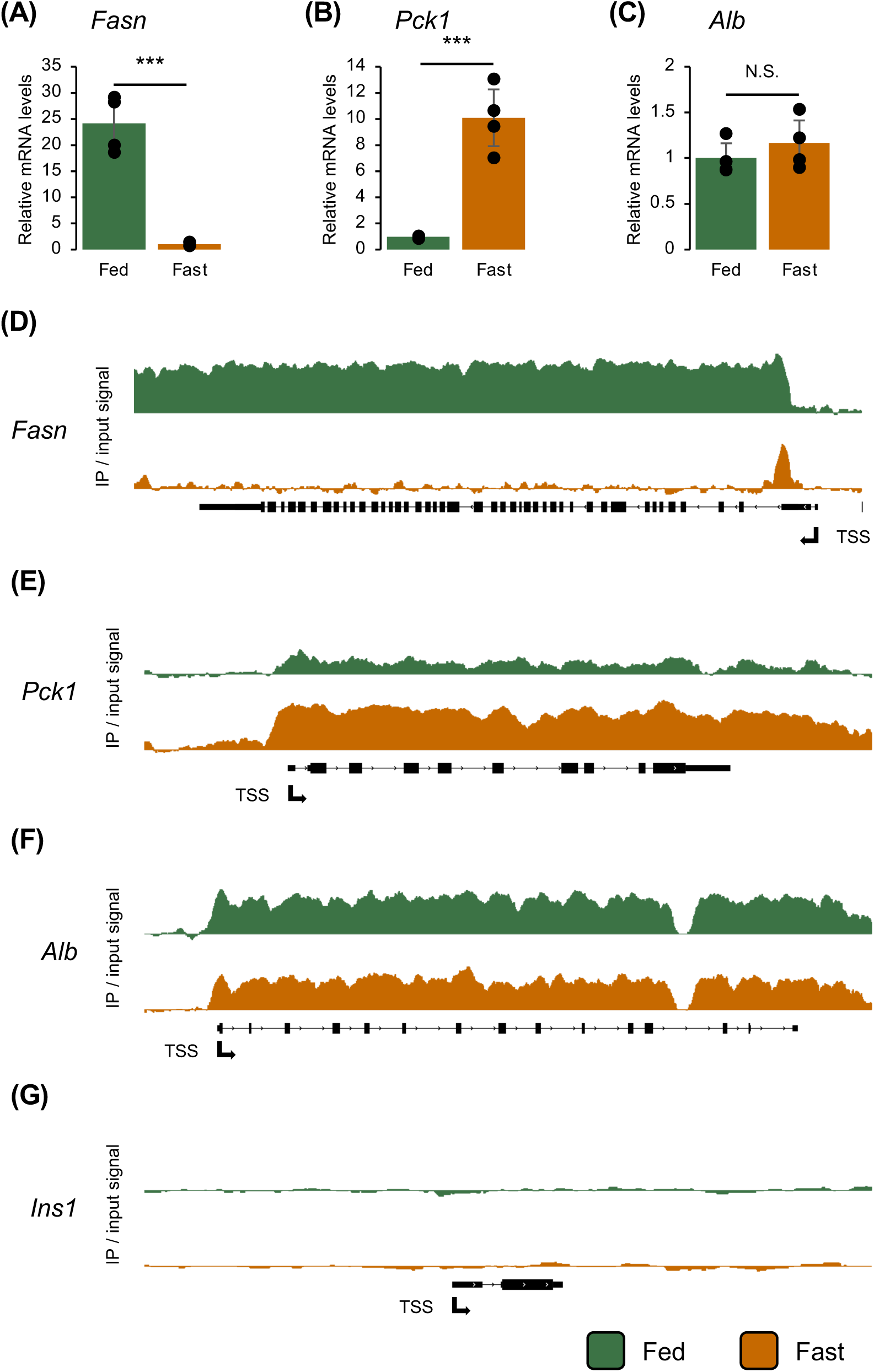
*Pck1* is transcribed in hepatocytes in the fed state. (A) Relative mRNA levels of *Fasn* in fed/fasted total hepatocytes. (B) Relative mRNA levels of *Pck1* in fed/fasted total hepatocytes. (C) Relative mRNA levels of *Alb* in fed/fasted total hepatocytes. (D) RNAP II ChIP of *Fasn* in fed/fasted total hepatocytes. (E) RNAP II ChIP of *Pck1* in fed/fasted total hepatocytes. (F) RNAP II ChIP of *Alb* in fed/fasted total hepatocytes. (G) RNAP II ChIP of *Ins1* in fed/fasted total hepatocytes.

The apparent transcription of gluconeogenic genes in the fed state could occur in all hepatocytes or be restricted to a subset of hepatocytes. To explore these possibilities, we isolated pericentral and periportal hepatocytes from fed and 16 h fasted mice using a digitonin ablation method that we modified for application in mice^5,14,15^. The isolated pericentral (PC) and periportal (PP) hepatocytes were then subjected to RNAPII ChIP-seq analyses. To confirm the relative purity of separation we examined the mRNA expression levels of *Cyp2e1*, an established PC hepatocyte marker, and *Cyp2f2*, an established PP marker^3,5,16^. As shown in Figure S2A there was a ∼ 6 to 9-fold enrichment of *Cyp2e1* in the PC hepatocytes and an ∼ 4 to 6-fold enrichment of *Cyp2f2* in the PP hepatocytes (Fig. S2B).

RNAPII ChIP-seq demonstrated that *Fasn* gene was transcriptionally active in both the PC and PP hepatocytes in the fed state, although somewhat greater in the PP hepatocytes (Fig. S2C). The *Fasn* gene occupancy of RNAPII in both PC and PP hepatocytes is consistent with the mRNA expression of lipogenic genes levels across the liver lobule^5^. In any case, in the fasted state there was essentially no RNAPII occupancy of *Fasn* gene in either PC or PP hepatocytes with the proximal promoter pausing peak readily observable in both zones. As expected in the fasted state, RNAPII occupancy of the *Pck1* gene was high in both the PC and PP hepatocytes (Fig. S2D), consistent with *Pck1* mRNA expression that is also expressed in both PC and PP hepatocytes following long-term fasting^5^. However, in contrast to *Fasn* gene, in the fed “off-state” *Pck1* gene still displayed substantial RNAPII occupancy in the PP hepatocytes but nearly background levels in the PC hepatocytes (Fig. S2D). As controls, RNAPII occupancy of *Alb* was neither zonated nor responsive to fasting/feeding (Fig. S2E) and the RNAPII tracks for *Ins1* was essentially background (Fig. S2F). These data indicate that the basal (fed state) transcription and expression of gluconeogenic genes occur predominantly in PP hepatocytes.

### Lipogenic and gluconeogenic genes are co-expressed in a subset of periportal hepatocytes

We next addressed whether all PP hepatocytes or a subset of PP hepatocytes were responsible for the basal expression of gluconeogenic genes in the fed state. To address this, we first utilized quantitative targeted single cell RNA-seq (targeted scRNA-seq) using the 10x genomic platform for a selective subset of metabolic genes^5^. Cell ranger and Loupe Browser were used for analyses for the quantitative targeted scRNA-seq to establish UMAP cluster maps of the hepatocytes in the fully fed and fasted states (Fig. 2A). To identify the position of the PC and PP hepatocytes (Fig. 2B) we took advantage of previous studies that have characterized the localization of specific zonation markers^3,5,16^. As examples, the PC markers *Cyp2e1* and *Gulo* (Fig. 2C) were highly colocalized to one region whereas the PP markers *Cyp2f2* and *Hsd17b13* (Fig. 2D) were highly colocalized to another distinct region. As these genes are typically expressed as a gradient across the lobule, (with the notable exception of *Glul* that is highly restricted to hepatocytes in one to two layers around the central vein), the other lower expressing cells likely represent the mid-zone hepatocytes^1,5^. As expected, in the fully fed state, the *de novo* lipogenic gene *Fasn* was found in both the PC and PP hepatocytes but following fasting was essentially absent from all the hepatocytes (Fig. 2E). On the other hand, the gluconeogenic gene *Pck1* was expressed in both the PC and PP hepatocyte clusters in the fasted state, but also showed expression in a subset of PP, but not PC, hepatocytes in the fed state (Fig. 2F). Alignment of the UMAPs for the hepatocytes that co-express both *Fasn* and *Pck1* mRNA in the fed state indicated they account for 7% of the periportal hepatocytes (Fig. 2G). In addition, comparison of other core *de novo* lipogenic genes (*Acly*, *Acaca*, and *Elvol6*) with the gluconeogenic gene *G6pc* also demonstrated that approximately 19% of PP hepatocytes simultaneously co-expressed these mRNAs (Fig. S3A-E).

**Figure 2.**
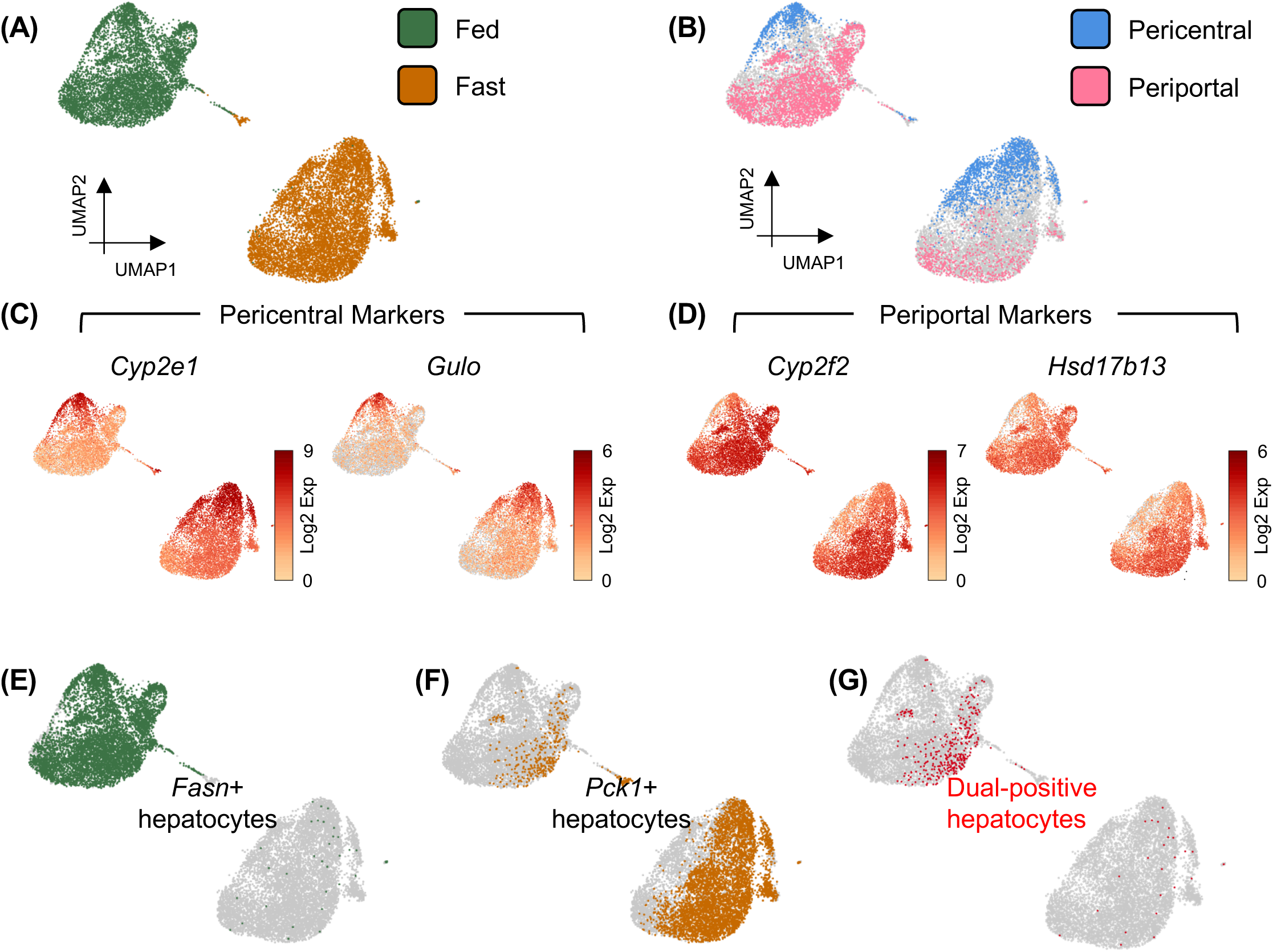
*Fasn* and *Pck1* are co-expressed in a subset of periportal hepatocytes. (A) UMAP visualizations of the targeted scRNA-seq based on feeding conditions. (B) UMAP visualizations based on zonation. (C) UMAP of PC zonation markers *Cyp2e1* and *Gulo.* (D) UMAP of PP zonation markers *Cyp2f2* and *Hsd17b13.* (E) UMAP of *Fasn* positive hepatocytes. (F) UMAP of *Pck1* positive hepatocytes (G) UMAP of dual-positive hepatocytes.

### *Fasn* and *Pck1* are actively co-transcribed in a subset of periportal hepatocytes

To confirm the scRNA-seq identification of a periportal hepatocyte subset that simultaneously expresses both gluconeogenic and *de novo* lipogenic genes, we utilized Hybridization Chain Reaction RNA Fluorescence *in Situ* Hybridization (HCR RNA-FISH) to simultaneously visualize the transcription sites (TS) of the *Fasn* and *Pck1* genes. Simultaneous glutamine synthetase RNA-FISH was used to identify the PC region (Figure 3, *Glul* not shown)^1,5^. In the fed state *Fasn* TS were readily observed throughout the lobule but fewer *Pck1* TS were detected in a subset of the PP but not PC hepatocytes (Fig. 3A, Fig. S4A and S4B). Magnified images clearly demonstrated the presence of both *Fasn* and *Pck1* TS simultaneously in the same nucleus in a subset of the PP hepatocytes (Fig. 3B). Quantification indicated that 24 % of the periportal hepatocytes and 1 % of the pericentral hepatocytes displayed TS for both *Fasn* and *Pck1* in the fed state (Fig. 3C). On the other hand, in the fasted state, *Pck1* TS was readily detected throughout the lobule but with essentially no *Fasn* TS (Fig. 3D, Fig. S4C and S4D). Magnified images of TS in the fasted state are shown in Fig. 3E. Quantification indicated that hepatocytes that were dual-positive of TS were not detected (Fig. 3F). These data directly demonstrate that the subset of PP hepatocytes actively transcribed both lipogenic (*Fasn*) and gluconeogenic (*Pck1*) genes at the same time.

**Figure 3.**
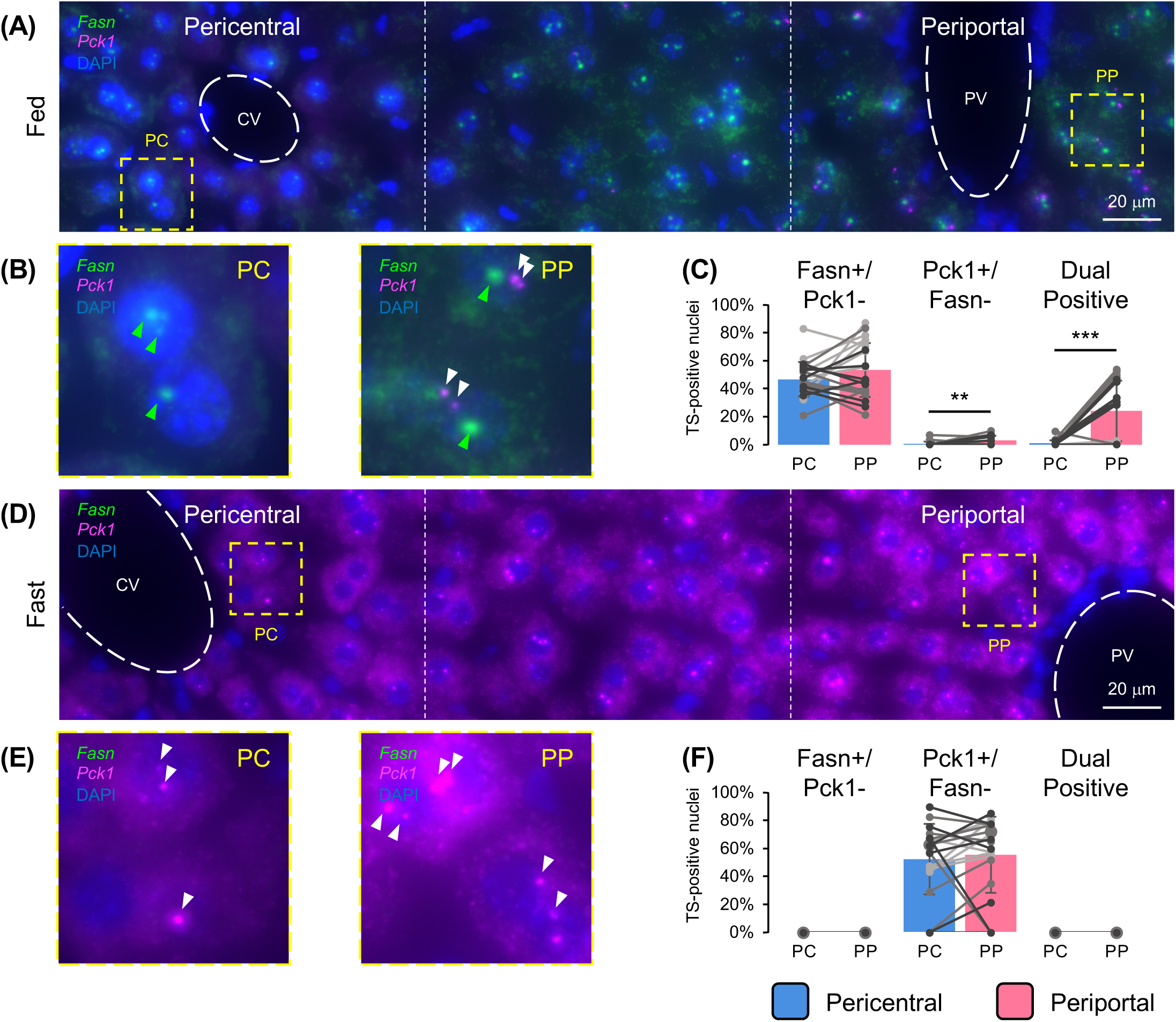
*Fasn* and *Pck1* are actively co-transcribed in a subset of periportal hepatocytes. (A) HCR RNA-FISH image of *Pck1* (magenta) and *Fasn* (green) with DAPI-stained nuclei (blue) in the fed state (representative image shown). Lobules show the central vein (CV) on the left and the portal vein (PV) on the right. (B) Magnified images of nuclei in box in panel (A). Arrows show transcription sites (TSs) (green arrow=*Fasn*, white arrow=*Pck1*, individual channel images of *Pck1* and *Fasn* are shown in Figure S4A and S4B). (C) Quantitative analysis of TSs in the fed state. Lines connect TS quantification between PC and PP within a lobule (N=21 lobules). Data are from 3 individual mice and are represented with different colors. Mean ± s.d. is indicated. *P*-value was calculated by paired *t*-test, N.S. if not indicated. 627 PC nuclei, 632 PP nuclei were analyzed, Scale bar, 20 um. (D) HCR-RNA FISH image of *Pck1* (magenta) and *Fasn* (green) with DAPI-stained nuclei (blue) in the fasted state (representative image shown). Lobules show the central vein (CV) on the left and the portal vein (PV) on the right. (E) Magnified images of nuclei in box in panel (D). White arrows highlight Pck1 TS (individual channel images of *Pck1* and *Fasn* are shown in Figure S4C and S4D). (F) Quantitative analysis of TS in the fasted state. Lines connect TS quantification between PC and PP within a lobule (N=21 lobules). Data are from 3 individual mice and are represented with different colors. Mean ± s.d. is indicated. *P*-value was calculated by paired *t*-test, N.S. if not indicated. 705 PC nuclei, 822 PP nuclei were analyzed, Scale bar, 20 um.

### Periportal hepatocytes have basal pyruvate-to-glucose flux in the fed state

Gene transcription and mRNA expression does not necessarily indicate functional activity; we therefore directly determined the relative *de novo* lipogenesis and gluconeogenesis activities by quantifying the fractional incorporation of ^2^H_2_O into palmitate and [2,3-^13^C_2_] pyruvate into glucose (Fig. 4). As expected, the fractional amount of ^2^H-palmitate in total hepatocytes from the fed state was substantially higher (16-fold) compared to hepatocytes from the fasted state (Fig. 4A). The decomposition of this overall palmitate enrichment is shown as the increased fed vs. fasted isotopologue enrichment distribution (Fig. 4B). This large increase appears to result from the very low fractional synthesis in the fasted state. Conversely, the average ^13^C incorporated per molecule of glucose synthesized was higher (2.4-fold) in the hepatocytes from the fasted state compared to the fed state (Fig. 4C). The decomposition of the isotopologue distribution of glucose is shown in Figure 4D with the majority of glucose isotopologues synthesized as either M2- or M4-glucose. To address the potential zonation of both glucose and palmitate synthesis, we next examined the relative synthesis rates in isolated PC and PP hepatocytes.

**Figure 4.**
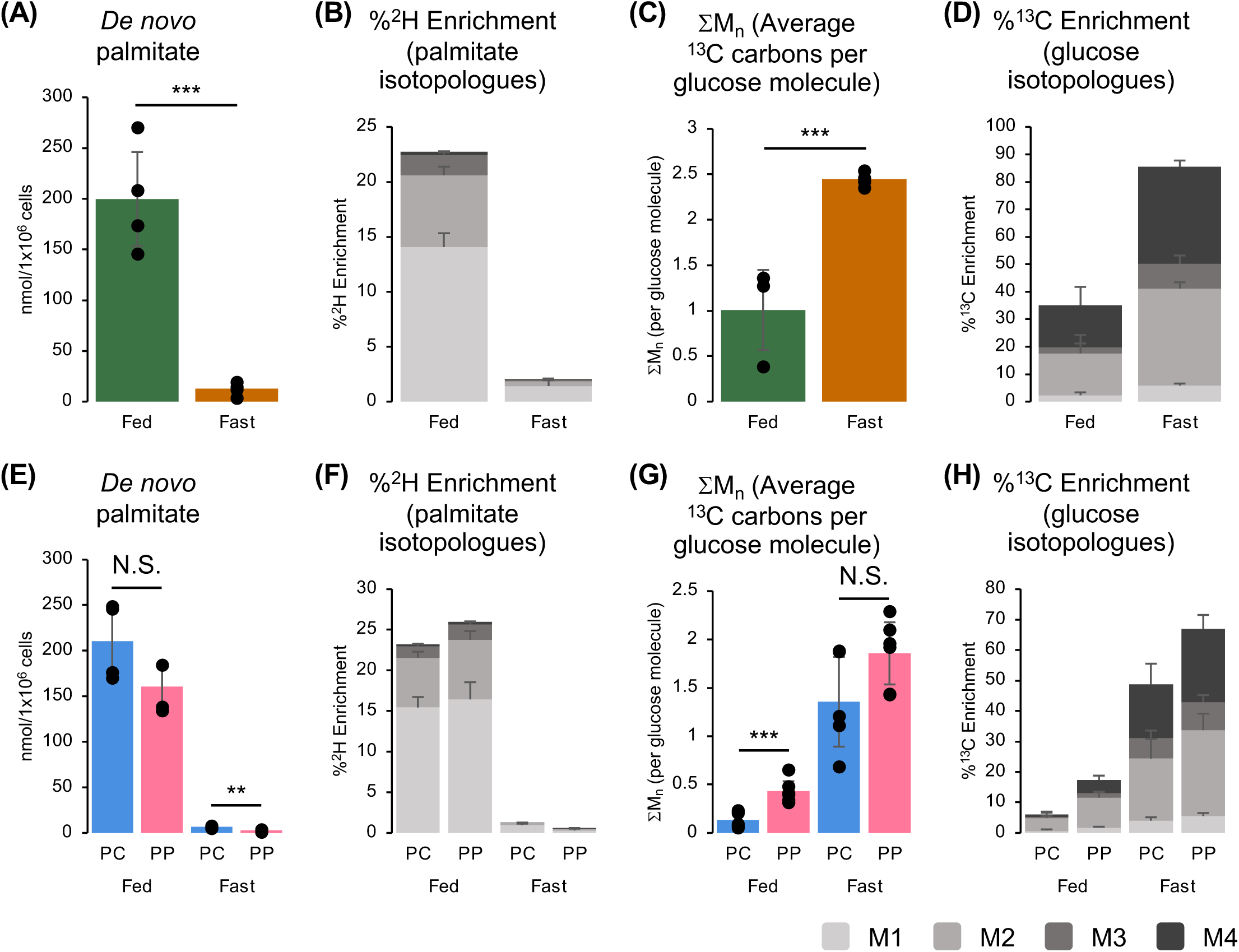
Periportal hepatocytes have basal pyruvate-to-glucose flux in the fed state. (A) *De novo* palmitate measurements using deuterated water in total hepatocytes from fed (n=4) and fasted (n=4) mice. Data are presented as mean values; error bars, s.d. Comparison between groups was made with unpaired, two-tailed Student’s *t*-test (***P < 0.001). (B) Fed and fast palmitate isotopologues derived from ^2^H_2_O in total hepatocytes. (C) ΣM_n_ (average ^13^C carbons per molecule) of glucose derived from [2,3-^13^C_2_]-pyruvate in total hepatocytes from fed (n=3) and fasted (n=5) mice. Data are presented as mean values; error bars, s.d. Comparison between groups was made with unpaired, two-tailed Student’s *t*-test (***P < 0.001). (D) Fed and fast glucose isotopologues derived from [2,3-^13^C_2_]-pyruvate in total hepatocytes. (E) De novo palmitate measurements using deuterated water in PC/PP hepatocytes from fed (n=4) and fasted (n=4) mice. Data are presented as mean values; error bars, s.d. Comparison between groups was made with unpaired, two-tailed Student’s *t*-test (**P < 0.01). (F) Fed and fast palmitate isotopologues derived from ^2^H_2_O in PC/PP hepatocytes. (G) ΣM_n_ (average ^13^C carbons per molecule) of glucose derived from [2,3-^13^C_2_]-pyruvate in pericentral (PC) and periportal (PP) hepatocytes from fed and fasted mice (n=5 for PC, n=6 for FastPP, n=7 for FedPP). Data are presented as mean values; error bars, s.d. Comparison between groups was made with unpaired, two-tailed Student’s *t*-test (***P < 0.001). (H) Fed and fast glucose isotopologues derived from [2,3-^13^C_2_]-pyruvate in PC/PP hepatocytes.

Consistent with *Fasn* gene expression, *de novo* lipogenesis activity is relatively high in the fully fed state (Fig. 4E) with very low activity in the fasted state and exhibits similar levels between PC and PP hepatocytes. The fold stimulation from the fasted “off” state and the active fully fed “on” state is approximately 30-fold in PC and 55-fold in PP hepatocytes. The isotopologue distribution of palmitate is shown in Figure 4F. In contrast, in the fed state there is a low level of gluconeogenesis from isolated PC hepatocytes, but it is several-fold higher in the PP hepatocytes (Fig. 4G). In the fasted state, both PC and PP hepatocytes display a higher rate of gluconeogenesis. Importantly, the PC hepatocytes increase their gluconeogenic activity ∼10-fold, whereas the PP hepatocytes only increase their gluconeogenic activity ∼4-fold due to the higher basal activity (Fig. 4G). The isotopologue distribution of glucose is shown in Figure 4H, which again shows mainly the formation of M2- and M4-glucose. These data demonstrate that in the fed state, PP hepatocytes display a significant degree of gluconeogenic activity, consistent with the PP expression of gluconeogenic genes in the fed state. On the other hand, PC hepatocytes exhibit a metabolic plasticity which enables them to become equally gluconeogenic in the fed to fasted transition.

### Dual-positive hepatocytes are insulin resistant in suppression of *Pck1*

Since the fed state is a normal physiologically anabolic state primarily driven by insulin, the expression of *Pck1* in the fed state suggests that this subset of PP hepatocytes was less responsive to insulin (i.e., insulin resistant), at least to insulin suppression of *Pck1* gene expression and gluconeogenic activity. To address this, we established a quantitative assay to isolate these dual expression (*Fasn* and *Pck1*) hepatocytes using PrimeFlow that is a combination of *in situ* hybridization coupled with flow cytometry^17^. As shown in Figure S5A hepatocytes are relatively large cells with a broad forward (FSC-A) and side scatter (SSC-A) parameters. Primary hepatocytes also display background fluorescence in the APC-Cy7 and mCherry channels, which were used for the *Pck1* and *Fasn* fluorescent mRNA probes (Fig. S5B). Nevertheless, in the fed state there is a substantial increase in the *Fasn* signal, resulting in 75 % of hepatocytes positive for *Fasn* (Fig. S5C). Similarly, in the fasted state there is a substantial increase in the *Pck1* signal with 71 % of the hepatocytes positive for *Pck1* (Fig. S5D). Having established this gating range, when we co-label the fed state hepatocytes with both the *Pck1* and *Fasn* probes and identified that approximately 12% of the hepatocytes were dual-positive (Fig. S5E). With a quantitative method in hand, we determined the number of dual *Pck1*/*Fasn* positive hepatocytes relative to the single positive and dual negative hepatocytes and performed euglycemic-hyperinsulinemic clamps in the 24-hour fasted state and the fed state at various insulin infusion rates. We also determined the relative suppression of the *Pck1* and *G6pc* mRNA in total hepatocytes. Under these conditions, the fasted *Pck1* and *G6pc* mRNA expression levels significantly decreased at 0.3 mU/kg/min (Fig. S5F and S5G). In contrast, fed *Pck1* mRNA in the dual-positive hepatocytes was not suppressed until 2 mU/kg/min of infused insulin (Fig. S5H). These data directly demonstrate that the *Fasn*/*Pck1* dual-positive hepatocytes are transcriptionally insulin resistant compared to the bulk liver hepatocytes.

Together these findings demonstrate that a subset of PP hepatocytes simultaneously express both gluconeogenic and lipogenic genes in the fed state (Figs. 2, 3, S3 and S4), and that isolated PP hepatocytes can also display both gluconeogenic and *de novo* lipogenic activities in the fed state (Fig. 4). However, this data does not directly demonstrate that individual dual-positive hepatocytes are in fact dual functional. To test for this possibility, we employed single cell spatial metabolic imaging to simultaneously assess both gluconeogenic and *de novo* lipogenic activities within individual hepatocytes

### A subset of liver regions has higher ATP turnover in the fed state

Depending upon the substrate, gluconeogenesis requires 6-10 ATP molecules per molecule of glucose generated. Similarly, synthesis of palmitate from acetyl-CoA requires 7 ATP and 14 NAPDH molecules. Thus, if a single hepatocyte is simultaneously engaged in both gluconeogenesis and *de novo* lipogenesis, it can be hypothesized that there must necessarily be a high rate of ATP turnover compared to the other bulk hepatocytes. During the process of energetic addition of phosphates to nucleotides or metabolites, there is an exchange of the oxygen in the phosphate with the oxygen in cell water^18–21^. ATP turnover reflects the percentage of ^16^O in ATP replaced by ^18^O after the *in vivo* administration of ^18^O-labeled water, reflecting whole cell energetics, as the exchange of ^18^O in high energy phosphates includes ATP synthesis and ATP hydrolysis, and the ^18^O can also be exchanged with creatine kinase and adenylate kinase catalyzed phosphotransfer^22,23^ (Table S1). Thus, we treated mice with or without ^18^O labeled water *in vivo* coupled with spatial imaging by DESI-Cyclic IMS TOF. Figure 5 shows liver sections from the fed and 24 h fasted state that were either untreated (unlabeled) or treated with ^18^O labeled water as we previously reported ^23^. Mass spectrometry imaging was done at 60×60 µm for the mass of normally abundant m0 (unlabeled) AMP in the negative ion mode. As mouse hepatocytes are polygonal and are nominally 20-30 µm in diameter, this resolution likely reflects the combination of several cells^24^. Regardless, the relative amount of the m0 AMP indicated a relatively low signal in the fed state but was markedly increased in the fasted state, indicating a greater turnover of AMP in the fasted state (Fig. 5A). In contrast, imaging for ^18^O-labeled AMP (m0+2 AMP) demonstrated an increased signal from background in both the fasted and fed states, from the mice treated with ^18^O labeled water compared to control mice livers (Fig. 5B). More importantly, the liver cross-section in the fasted state showed increased intensities suggesting increased turnover relative to the fed state (Fig. 5B). In contrast, m0+2 ADP fasted and fed intensities were similar over each slice, suggesting turnover was similar between the fed and fasted state. The increase of m0+2 ATP intensities over the liver cross-section suggested turnover was increased in the fed state compared to the fasted state, which would correspond to an increase in ATP generation in the fed state (Fig. 5C). As these AMP, ADP and ATP measurements were done on the same slice, m0+2 ADP and m0+2 ATP pixel densities can be compared to m0+2 AMP in fed state with multiple sites of overlap between m0+2 AMP, ADP and ATP (Fig. 5D). These results revealed several discrete pixels that clearly overlapped between m0+2 AMP (red), ADP (blue) and ATP (green) (Fig. 5D). The white areas indicate pixel regions with strong overlap for all three signals. These data strongly indicate that a small subset of liver regions have higher rates of ATP, ADP and AMP turnover in the fed state, indicating increased energetics, which might be expected for Dual-Modal hepatocytes.

**Figure 5.**
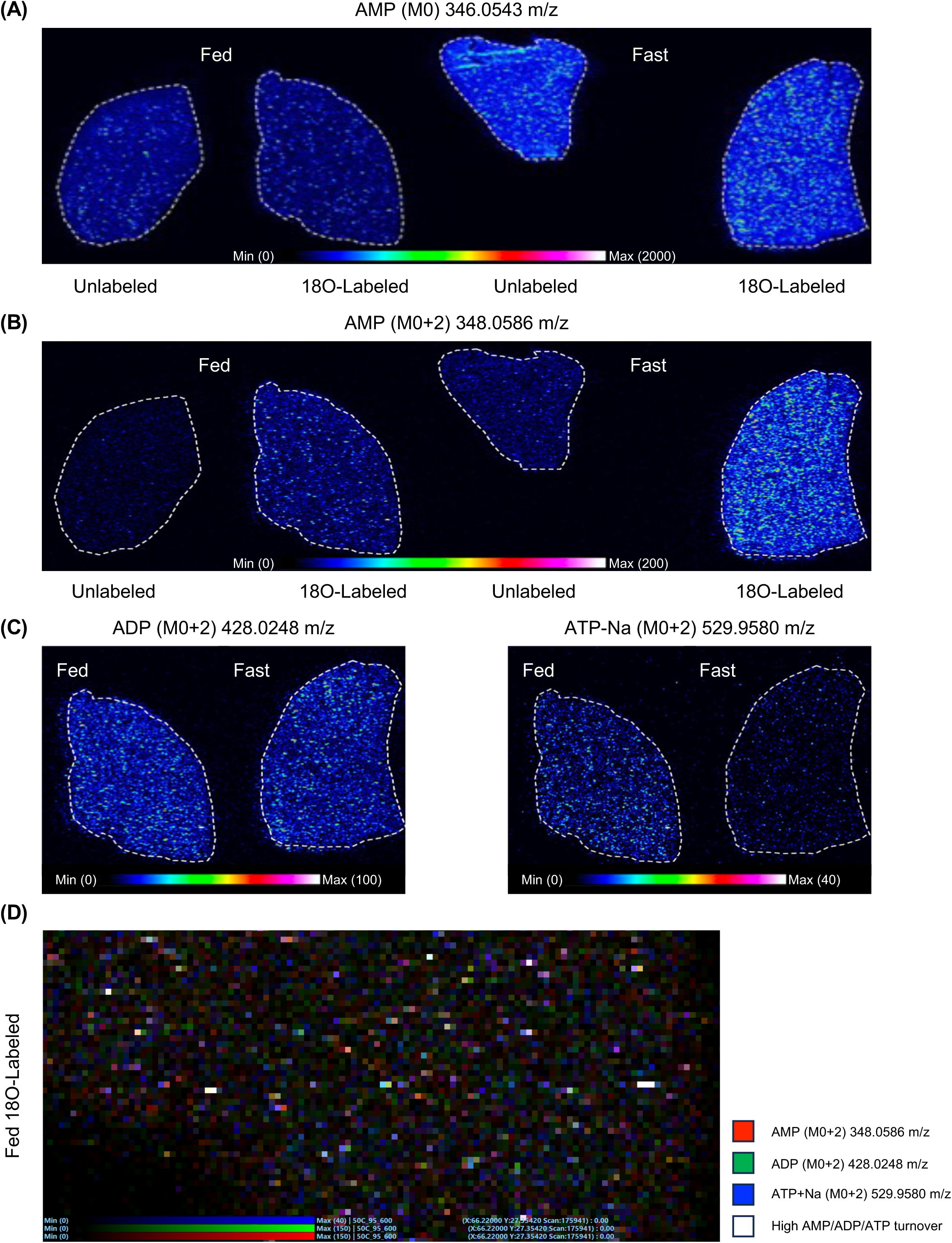
A subset of liver regions has higher ATP turnover in the fed state. (A) Representative DESI image of unlabeled AMP (M0) from unlabeled and 18O labeled water intraperitoneal+gavage mice in the fed and fasted state. (B) Representative DESI image of labeled AMP (m0+2) from unlabeled and 18O labeled water intraperitoneal+gavage mice in the fed and fasted state. (C) Representative DESI image of labeled ADP (m0+2) (left) and ATP-Na (m0+2) (right) from 18O labeled water intraperitoneal+gavage mice in the fed and fasted state. (D) Representative DESI image of labeled AMP (m0+2, red), ADP (m0+2, blue), and ATP-Na (m0+2, green) merged as one image. The white areas indicate pixel regions with strong overlap for all three signals.

### A subset of hepatocytes is lipogenic and gluconeogenic

To more specifically address whether an individual hepatocyte can simultaneously undergo both *de novo* lipogenesis and gluconeogenesis, we next utilized metabolic spatial imaging to identify the formation of m0+4 palmitate (*de novo* lipogenesis) and m0+3 glucose-6-phosphate (gluconeogenesis) from a liver section of mice *in vivo* fed with U[^13^C]-glucose. m0+3 glucose-6-phosphate (3 labeled carbons) can only be formed from U[^13^C]-glucose (6 labeled carbons) when the m0+6 glucose is metabolized to a m0+3 triose. The triose then goes up the gluconeogenic pathway to glucose or first down the glycolytic pathway into the TCA cycle and then back up the glycolytic pathway to form glucose (Fig. 6A). On the other hand, labeled palmitate can only be formed if the U[^13^C]-glucose becomes m0+3 pyruvate and then enters the TCA cycle through pyruvate dehydrogenase to generate the 2-carbon labeled acetyl-CoA. Subsequent metabolism and addition of these 2-carbon labeled units for fatty acid synthesis results in the formation of m0+2, m0+4 etc. labeled palmitate (Fig. 6A). The dual-imaging of m0+4 palmitic acid (red) and m0+3 glucose-3-phosphate (blue) by DESI MSI at 25 x 25 µm demonstrated the presence of pixels that were dual positive for both (pink, indicated by arrows) (Fig. 6B). The same experiment was performed with MALDI MSI in a lower magnification (50 x 50 µm). The ingestion of U[^13^C]-glucose in the fed state resulted in the appearance of m0+4 palmitate acid (red) and m0+3 glucose-6-phosphate (blue) throughout the liver section whereas there was no detectable background labeling when ingestion of unlabeled glucose in a parallel mouse (Fig. 6C). A higher magnification of this MALDI image showed the presence of pixels that were dual positive for both m0+4 palmitic acid and m0+3 glucose-6-phosphate (pink, indicated by arrows) (Fig. 6D). Since this resolution does not provide single cell information^24^, we next performed mass spectrometry spatial imaging at 10 µm resolution for m0+4 palmitic acid (red) and m0+3 glucose-6-phosphate (blue) that we co-registered with H&E staining (Fig. 6E and F). As is apparent, in this imaging each pixel is smaller than the hepatocytes and with a high probability it represents a single hepatocyte (Fig. 6E and F). Consistent with the lower resolution image, individual hepatocytes mostly display *de novo* lipogenesis in the fed state (m0+2 palmitic acid positive, red) with fewer hepatocytes positive for m0+3 glucose-6-phopshate (blue). However again, there were several pixels in the field that are dual positive (pink), consistent with the overlap seen with ATP, ADP and AMP turnover (Fig. 5) and the dual-positive hepatocytes present as measured by both scRNA-seq and PrimeFlow. As these data confirm the metabolic functional duality (simultaneous gluconeogenic and *de novo* lipogenic activities), we have termed these as Dual-Modal hepatocytes.

**Figure 6.**
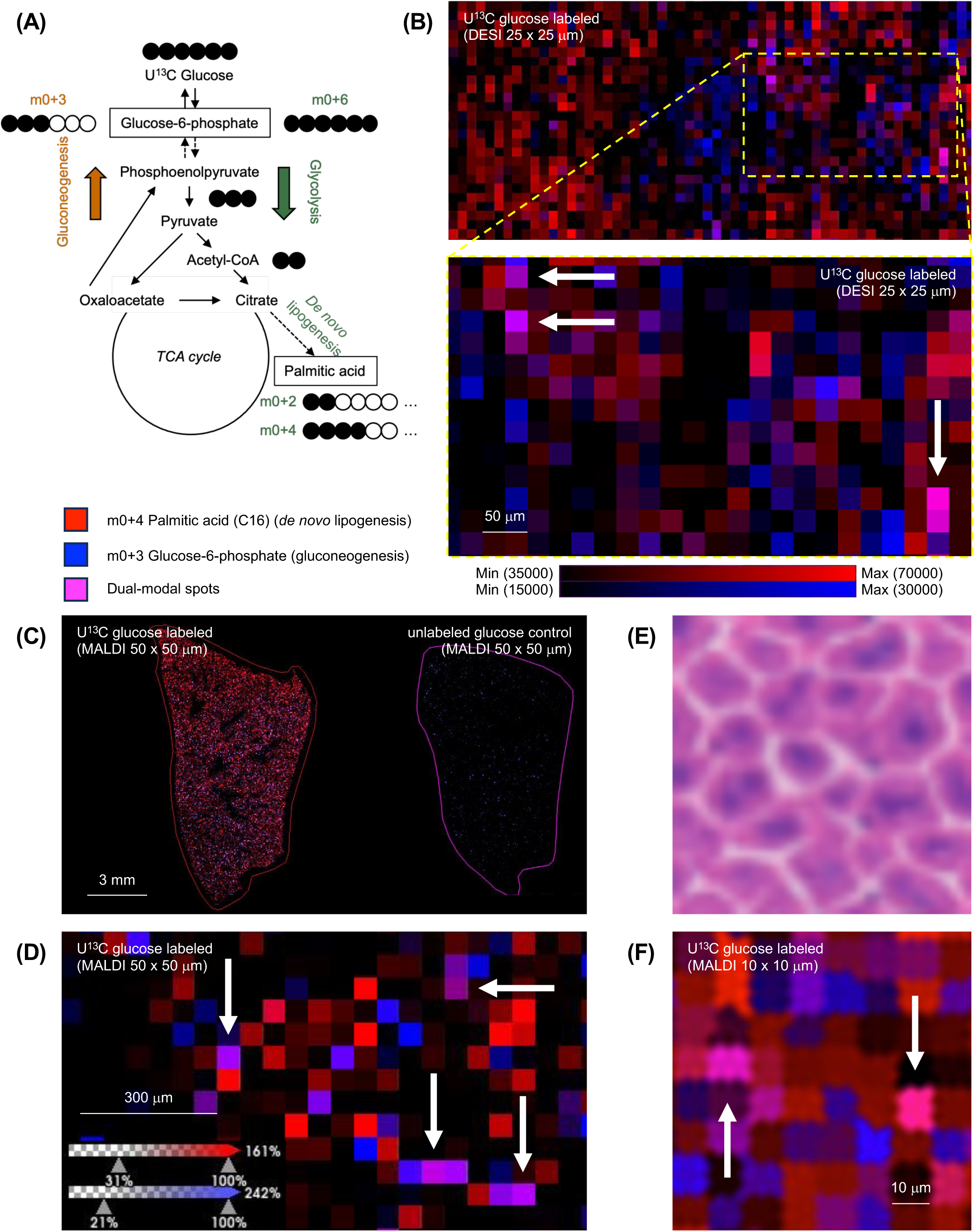
A subset of hepatocytes is lipogenic and gluconeogenic. (A) Scheme of detecting Dual-modal (DM) spots with U^13^C glucose. (B) 25 x 25 µm DESI image of U[^13^C]-glucose labeled m0+4 Palmitic acid (red) and m0+3 Glucose-6-phosphate (blue); pink represent the DM spots. White arrows highlight the DM spots. (C) Low magnification MALDI image of U[^13^C]-glucose labeled m0+4 Palmitic acid (red) and m0+3 Glucose-6-phosphate (blue); pink represent the DM spots, unlabeled glucose section as control. Each pixel is 50 µm. (D) High magnification MALDI image of U[^13^C]-glucose labeled m0+4 Palmitic acid (red) and m0+3 Glucose-6-phosphate (blue); pink represent the DM spots. White arrows highlight the DM spots. Each pixel is 50 µm. (E) High magnification of H&E staining. (F) High magnification MALDI image of U[^13^C]-glucose labeled m0+2 Palmitic acid (red) and m0+3 Glucose-6-phosphate (blue); pink represent the DM hepatocytes. White arrows highlight the DM hepatocytes. Each pixel is 10 µm.

### Humanized mouse liver has dual-positive spots

Having established the presence of hepatocyte subsets that simultaneously express both *Fasn* and *Pck1* mRNA in the fed state of mouse liver, we addressed whether dual-positive hepatocytes are also present in human liver hepatocytes. As patient surgical interventions can only be performed in the fasted state, it is not possible to obtain liver specimens or biopsy samples from humans in the fed state. Thus, we utilized a humanized mouse model in which human hepatocytes replace greater than 95 % of the mouse hepatocytes and performed 10x Genomics spatial transcriptomics Visium analyses on liver sections^25^.Similar to the mice data, RT-qPCR for human *Fasn* showed relatively low expression in the fasted state with induction in the fed state (Fig. S6A). Additionally, the human *Fasn* PCR primers were unable to detect *Fasn* mRNA in wild-type mouse liver, indicating no cross-reactivity. Similarly, the human *pck1* PCR primers did not cross-react with mouse *Pck1* but clearly displayed the fasted state induction of *Pck1* (Fig. S6B). As the Visium protocol requires a prior fixation step, we also confirmed that this fixation step did not disrupt the integrity of mRNA detection by RT-qPCR (Fig. S6C and D). Relative differences in the expression of the human PC marker (*CYP1A2*) and PP marker (*HSD17B13*) as shown in the violin plots (Fig. S6E and S6F) enabled the identification of PC spots (blue) and PP spots (red) (Fig. S6G). The violin plots for *Fasn* (Fig. S6H) and *Pck1* (Fig. S6I) display the expected zonation, in particular the *Pck1* mRNA in the PP hepatocytes are elevated compared to those of the PC hepatocytes. The distribution of the *Fasn*/*Pck1* dual-positive spots is marked in yellow with the H&E staining of the section (Fig. S6J) and with a magnified region show in Figure S6K. Quantification showed 9% of the PP hepatocytes were *Fasn*/*Pck1* dual-positive (Fig. S6L). These results suggest Dual-Modal hepatocytes presumable exist in human hepatocytes as well.

### Dual-positive spots increase in high-fat diet fed mice liver

Lastly, having established the Dual-Modal (*Fasn*/*Pck1* positive) hepatocytes that are naturally insulin resistant (Fig. S4) and are gluconeogenic in the fed state, we speculated that in states of insulin resistance the number of these hepatocytes might increase. To test this hypothesis, we determined the number of Dual-Modal hepatocytes in mice fed a normal chow diet (NCD) or a high-fat diet for 3 months (3MHFD) and used the Visium platform for spatial single dot analyses. t-SNE plots showed separation between NCD and 3MHFD (Fig. S7A). Graph-based clustering showed zonation patterns in both NCD and 3MHFD (Fig. S7B, S7C). Based on analyses of several zonation markers such as *Cyp2e1* and *Glul* for pericentral (Fig. S7D) and *Cyp2f2* and *Hsd17b13* for periportal (Fig. S7E), we identified the PC (pink) and PP (blue) spots in the t-SNE plot, showing clear separation (Fig. S7F). The PC and PP regions in comparison with histological H&E staining are shown in Figure 7A. The presence of lipid droplets in the HFD fed mice that are specifically localized to the PC hepatocytes is readily observed in the magnified region shown in Figure 7B, consistent with the known zonation of triglyceride accumulation^26^. The distribution of zonation markers as violin plots is shown in Figures 7C and 7D. As expected for the fed state, *Fasn* mRNA was expressed in both PC and PP hepatocytes in both the NCD and 3MHFD fed mice (Fig. S7G). Consistent with our previous findings, *Pck1* mRNA was also present in a subset of periportal hepatocytes (Fig. S7H) that overlapped with the *Fasn* dots (Fig. S7G). The overlap of *Fasn* and *Pck1* mRNA positive dots in the NCD and HFD fed mice is shown in Figures 7E. Quantification of the number of dots that were dual positive was approximately 4 % under NCD diet but following 3 months of HFD the number of dual positive dots increased to approximately 13 % (Fig. 7F). Moreover, these overlapping dots primarily occurred in the PP region (Fig. 7G). These data demonstrate that long term HFD results in an increase in the number of the Dual-Modal (*Fasn*/*Pck1* positive) hepatocytes.

**Figure 7.**
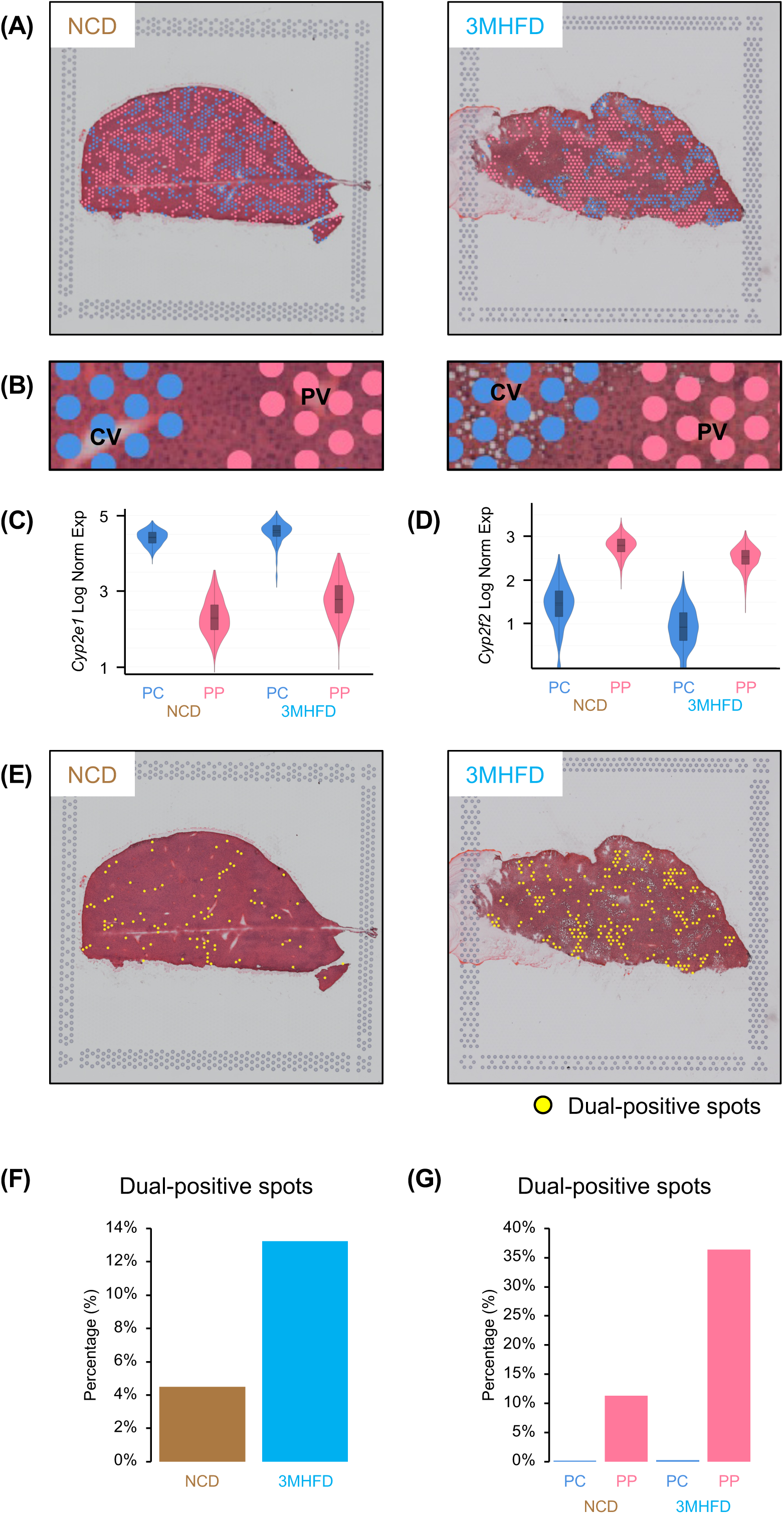
Dual-positive spots increase in high-fat diet fed mice liver. (A) Spatial transcriptomics image of Normal Chow Diet (NCD) fed mice and 3 Month High Fat Diet (3MHFD) fed mice with Pericentral (PC)/Periportal (PP) spots highlighted. (B) A representative lobule shown in a higher magnification with central vein (CV) on the left, portal vein (PV) on the right. Pink spots are PP, blue spots are PC. (C) Violin plot of *Cyp2e1* (pericentral (PC) marker) log normalized expression in PC/PP spots of NCD and 3MHFD fed mice. (D) Violin plot of *Cyp2f2* (periportal (PP) marker) log normalized expression in PC/PP spots of NCD and 3MHFD fed mice. (E) Distribution of DM spots in NCD and 3MHFD fed mice; yellow spots show DM spots. (F) Quantification of DM spots in NCD and 3MHFD fed mice. (G) Quantification of DM spots in PC/PP spots of NCD and 3MHFD fed mice.

## DISCUSSION

Gluconeogenesis is a fundamental metabolic process in which non-carbohydrate carbon substrates such as lactate, glycerol and amino acids are used to generate glucose^2,27^. In mammals’ gluconeogenesis is primarily performed in the hepatocyte, although kidney and intestine cells are capable of driving significant gluconeogenesis as well^27–29^. In general, gluconeogenesis is inhibited in the fed state when plasma glucose levels rise as a result of dietary absorption, and insulin secretion is stimulated. In the fasted state, dietary derived plasma glucose levels decline, reducing insulin secretion and stimulating glucagon and epinephrine release to activate hepatic gluconeogenesis and glucose release to restore and maintain a normal glycemic state. Two of the key gluconeogenic enzymes phosphoenolpyruvate carboxykinase (gene *Pck1*) and glucose 6-phosphate catalytic subunit (gene *G6pc*) are not thought to be allosterically regulated, but acutely controlled at the level of gene expression through upstream transcription factors that are activated by protein kinase A (i.e., CREB, HNF4α, FoxO1) signaling and suppressed by insulin signaling^27^.

Despite this generalized regulation of hepatic gluconeogenic gene expression and gluconeogenic activity in the fed and fasted states, previous studies assessing gluconeogenesis observed a basal level of glucose production from the liver even in the fed state^6,7^. In fact, examination of numerous hyperinsulinemic-euglycemic clamp data clearly shows that an insulin suppression of hepatic glucose production is never zero^30,31^. This has been typically ignored and assumed to simply represent a background signal. Our data suggest otherwise, as the mRNA expression and transcription of lipogenic genes are essentially zero in their respective off-state (fasted), whereas there is always a measurable level of gluconeogenic gene expression, transcriptional activation and gluconeogenic activity in their respective off-state (fed). This fed state mRNA expression of *Pck1* was confirmed by multiple, independent methodologies, including targeted quantitative single cell RNAseq, smFISH, HCR RNA-FISH and PrimeFlow. Moreover, PrimeFlow demonstrated the presence of both mRNA for *Fasn* and *Pck1* in the same cells and HCR RNA-FISH clearly visualized the same nuclei displaying transcription sites for both *Fasn* and *Pck1*. Each of these assays indicated a subset of periportal hepatocytes is dual *Fasn* and *Pck1* positive in the fed state.

Hepatocyte zonation is well established as a property of regionally distinct cellular functions localized across the liver lobule, and studies in the 1980’s using fasted rats reported that there was a 2 to 3-fold bias of gluconeogenesis in isolated PP hepatocytes relative to PC hepatocytes^14,32^. We also found an approximate 1.4-fold bias in gluconeogenic activity and *Pck1* gene expression in PP versus PC hepatocytes of fasted mice. However, analyses of gluconeogenic activity in the fed state, although absolutely less than in the fasted state, was approximately 3.2-fold greater in PP versus PC hepatocytes. In parallel, very few *Pck1* transcription sites and very low levels of *Pck1* mRNA or protein were detected in fed PC hepatocytes compared to PP hepatocytes. These data are consistent with a subset of PP hepatocytes being responsible for the basal (fed state) level of gluconeogenesis. As such, we refer to this hepatocyte subset as Dual-Modal (DM) hepatocytes.

The conversion of U[^13^C]-glucose to m0+2- and m0+4-palmitate is generally accepted as a direct measure of *de novo* lipogenesis that is readily detected by mass spectrometry imaging^33^. However, spatial mass spectrometry imaging cannot detect labeled glucose within a hepatocyte, as glucose is rapidly secreted. Instead, we measured hepatocyte content of m0+3-glucose-6-phosphate derived from ingested U[^13^C]-glucose. In order for U[^13^C]-glucose to generate m0+3-glucose-6-phosphate, the U[^13^C]-glucose must at minimum be converted by glycolysis or the pentose cycle into the triose pool, which in turn can either enter the TCA cycle (pyruvate->oxaloacetate->PEP->triose-P pool) for gluconeogenesis/glyceroneogenesis, or the trioses glycerol-3-phosphate and dihydroacetone phosphate recombine to form fructose-1,6-phosphate and subsequently glucose-6-phosphate. Regardless of whether recycled glucose is derived from the triose-P pool after traveling to the triose-P pool via glycolysis and the TCA cycle or the pentose cycle and/or via the TCA cycle, our finding of the generation of m0+3-glucose-6-phosphate from ingested U[^13^C]-glucose demonstrates that in the fed state, a subset of hepatocytes exhibits a simultaneous, bidirectional metabolic flow up and down the glycolytic pathway. Our finding that isolated PP hepatocytes in the fed state generate secreted glucose is consistent with a novel demonstration of gluconeogenesis in the fed state.

However, these data do not rule out other metabolic pathways for the generation of stable isotope labeled trioses. For example, the trioses could also contribute to glyceroneogenesis that in the fed state are needed for the generation of the backbone for triacylglycerol synthesis. Another more intriguing possibility is that the formation of m0+3 glucose-6-phosphate in the fed state may account for previous observations supporting an indirect pathway for glycogen synthesis^8–11^. After glycogen is depleted in the fasted state, it can be re-synthesized in the fed state by the direct pathway (dietary glucose converted to glucose-6-phosphate epimerized to glucose-1-phosphate that generates UDP-glucose for direct addition to glycogen^9,11^. Alternatively, glucose but also gluconeogenic substrates (i.e., amino acids) can be used to generate trioses by cycling down the glycolytic pathway into the TCA cycling and then metabolized via pyruvate carboxylase then PEPCK to generate triose-P which can then generate glucose-6-phosphate that is then used for the re-synthesis of glycogen^8–11^. Additional stable isotope flux analyses will be necessary to distinguish between these distinct routes of carbohydrate flux in this unique hepatocyte subpopulation, and which likely are not mutual exclusive.

While we observed that both normal mouse liver as well as humanized mouse livers express naturally insulin resistant, Dual-Modal hepatocytes, their functions in systemic physiology and pathophysiology remain to be determined. One possibility is that during ingestion of ketogenic diets, glucose, although lowered, still needs to be produced. In this case, Dual-Modal hepatocytes could provide a mechanism to maintain basal glycemic levels necessary for survival. Pathophysiologically, high-fat diet induced insulin resistance resulted in an increase in the number of Dual-Modal hepatocytes. This opens up the intriguing possibility that the change in proportion of this unique hepatocyte subpopulation may be an important component leading to hepatic insulin resistance.

### Limitations of the study

While we have identified a subset of hepatocytes termed Dual-Modal hepatocytes that could potentially explain the cause of hepatic insulin resistance based on an increased number of Dual-Modal hepatocytes in high-fat diet fed mice, it is important to recognize that these data do not distinguish between whether the number of Dual-Modal hepatocytes increase due to proliferation of the pre-existing Dual-Modal hepatocyte subpopulation and/or due to the development of insulin resistance in normal hepatocytes. Future lineage tracing studies will be necessary to assess these two possibilities. Also, this study does not molecularly address how an individual cell can apparently undergo gluconeogenesis and *de novo* lipogenesis at the same time, as these two metabolic pathways are generally assumed to be mutually exclusive with each other. The three most likely possibilities are i) futile cycling throughout the parallel glycolytic and gluconeogenic pathways for either periportal and/or pericentral hepatocytes (dependent on degree of fasted or refed states) thereby allowing substrates to flow in both directions; ii) relatively rapid oscillations between gluconeogenesis and *de novo* lipogenesis, perhaps operant in Dual-Modal hepatocytes, so that they are not actually occurring simultaneously; and iii) subcellular compartmentalization of metabolic flux such that different regions or mitochondria within an individual hepatocyte can undergo either gluconeogenesis or *de novo* lipogenesis.

## RESOURCE AVAILABLITY

### Lead contact

Further information and requests for resources and reagents should be directed to and will be fulfilled by the lead contact, Junichi Okada

### Materials availability

This study did not generate new unique reagents.

### Data and code availability

Raw ChIP and fastq files were deposited in the National Center for Biotechnology Information Gene Expression Omnibus database and are available under accession numbers: GSE315776, GSE263419 (previously published targeted scRNA-seq data) and GSE320372 (VISIUM). This study was conducted using only publicly available software; no custom code was used.

## ACKNOWLEDGEMENTS

We wish to acknowledge that the data presented in this manuscript was made possible through the expertise of The Albert Einstein College of Medicine Genomics, Animal Physiology, Flow Cytometry and Metabolomics Core Facilities. We are thankful to Floric Slimani for the smallFish package, Licheng Wu for euglycemic-hyperinsulinemic clamp studies, and Shiori Okada for data analysis. This study was supported by NIH grants DK020541, DK110063, DK069861, GM007491, CA013330, AI179019, AA031412, and the PhoenixBio R&D Grant.

## AUTHOR CONTRIBUTIONS

JO, AL, MH, AMX, RR, SVL, TS, GJS, CE, RCS, YQ and IJK all assisted in the experimental design, performance and analyses of the data presented, and writing of the manuscript. LL and KS assisted in the experimental design, bioinformatic analyses and editing of the manuscript. VLS, FY, MH, CE, KS and JEP supervised this project, experimental design, interpretation of data and writing the manuscript.

## METHODS

## KEY RESOURCES TABLE

| REAGENT or RESOURCE | SOURCE | IDENTIFIER |
| --- | --- | --- |
| Antibodies |  |  |
| RNAPII subunit B1 (Rpb1) NTD | Cell Signaling | 14958 |
| Chemicals, peptides, and recombinant proteins |  |  |
| Liberase | Sigma-Aldrich | 5401127001 |
| Digitonin | Sigma-Aldrich | D141-500MG |
| Percoll | Sigma-Aldrich | P4937-500ML |
| Critical commercial assays |  |  |
| ChIP DNA Clean & Concentrator kit | Zymo Research | D5205 |
| NEBNext® Ultra™ II DNA Library Prep Kit | New England Biolabs | E7645 |
| HCR RNA-FISH | Molecular Instruments | V3.0 (N/A) |
| PrimeFlow RNA Assay Kit | Invitrogen | 88-18005-204 |
| Deposited data |  |  |
| ChIP sequence | This paper, GEO | GSE315776 |
| Targeted scRNA-seq | Okada et al<sup>5</sup>, GEO | GSE263419 |
| Visium (spatial transcriptomics) | This paper, GEO | GSE320372 |
| Experimental models: Organisms/strains |  |  |
| C57Bl/6J | The Jackson Lab | 000664 |
| Oligonucleotides |  |  |
| Illumina® single Index Primers Set 1 | New England Biolabs | E7335 |
| Illumina® single Index Primers Set 2 | New England Biolabs | E7500 |
| See Table S2 for primer sequences | This paper | N/A |
| Software and algorithms |  |  |
| Trim Galore | Krueger et al | https://github.com/FelixKrueger/TrimGalore, V0.6.7 |
| Bowtie2 | Langmead et al<sup>34</sup> | V2.4.5 |
| Samtools | Li et al<sup>35</sup> | V1.6 |
| DeepTools | Ramirez et al<sup>36</sup> | V3.5.1 |
| Cell Ranger | 10x Genomics | V6.0.1 |
| Small Fish (Cellpose-SAM, big-fish) | Slimani et al | https://github.com/SmallFishGUI/small_fish_gui,<br>V2.1.10 |
| Space Ranger | 10x Genomics | V2.0.0, V3.1.0 |
| Other |  |  |
| PicoLab Rodent Diet 20 | LabDiet | 5053 |
| Rodent Diet with 60 kcal% Fat | Research Diets | D12492 |
| Modified OSD with Macronutrient Content Similar to LabDiet 5053 | Research Diets | D17011901px |

## EXPERIMENTAL MODEL AND STUDY PARTICIPANT DETAILS

### Animals

All animal studies were conducted in accordance with protocols approved by the Albert Einstein College of Medicine Institutional Animal Care and Use Committee (IACUC).

Unless otherwise indicated, experiments were performed using 10-14-week-old C57Bl/6J wild-type male mice (The Jackson Laboratory, 000664). Spatial transcriptomics studies were performed using 18-week-old mice. Animals were housed in an IACUC-approved facility under ambient temperature and humidity with a 12-hour light/dark cycle (lights turn on 7:00, lights turn off 19:00) and group housing. Mice had free access to water and laboratory mouse chow diet (LabDiet, 5053), unless otherwise specified. For high-fat diet studies, mice were fed a 60% fat diet (Research Diets, D12492) beginning at 6 weeks of age for 3 months.

To align feeding conditions, mice were trained for 3 days to a time-restricted feeding schedule by removing food at 17:00 and refeeding at 9:00 the following day (16-hour fasting and 8-hour feeding) as previously described^5,12^. On the day of the experiment, mice were provided food with drinking water supplemented with 20% sucrose (w/v) at 9:00. Fed mice were sacrificed at 13:00, corresponding to 4 hours of feeding. For fasted mice, food and sucrose-containing water were removed at 13:00, drinking water was replaced with regular water, and mice were fasted for 24 hours before sacrifice at 13:00 the following day. For pericentral/pericentral ChIP sequencing studies, mice were sacrificed after 16 hours of fasting. For high-fat diet studies (and control mice), food was removed at 7:00, refed at 19:00, and sacrificed at 23:00 (12-hour fasting and 4-hour feeding).

### Humanized liver chimeric mice

uPA SCID-based humanized liver chimeric mice were generated as previously described and maintained with vitamin C containing chow diet (modified LabDiet 5053 containing 130 ppm vitamin C)^25^. Humanized liver chimeric mice were subjected to a time-restricted feeding schedule modified to 10-hour fasting and 14-hour feeding.

## METHOD DETAILS

### Hepatocyte Isolation

Liver perfusion and hepatocyte isolation were performed under anesthesia as previously described^5^. For total hepatocyte isolation, a catheter was placed in the portal vein (PV) and the lower inferior vena cava (IVC) was cut for drainage. For zonated hepatocyte isolation, catheters were placed in both the PV and IVC, and the perfusion line was connected to the PV for periportal (PP) isolation of to the IVC for pericentral (PC) isolation. Livers were perfused with pre-warmed PBS (42 °C) at 5 mL/min, then switched to enzyme buffer solution (EBS, 15 mL) containing liberase (0.05 mg/mL) (Sigma-Aldrich, #5401127001). For zonated hepatocyte isolation, 833 μL of 5-10mM digitonin (Sigma-Aldrich, #D141-500MG) was injected via the unconnected catheter (IVC for PP, PV for PC), after which the vessel was cut, and perfusion was immediately resumed. Following perfusion, the liver was cut from the mouse and placed in cold DMEM (without glucose) or Krebs-Henseleit (KH) buffer, gently dissociated, and filtered through a 100 μm cell strainer. For liver hepatocyte purification, the cells were centrifuged at 40 x *g* for 5 min at 4 °C, followed by Percoll (Sigma-Aldrich, #P4937-500ML) solution (80 x *g* for 10 min at 4 °C), then washed by DMEM or KH buffer. Hepatocytes were counted and assessed for viability prior to downstream applications.

### ChIP sequencing

#### Fixation

Total hepatocyte (Fig. 1 and Fig. S1) and periportal/pericentral hepatocyte (Fig. S2) isolation was performed on fasted and fed mice as described above with the exception that all sample processing after liberase digestion was done using ice-cold DBPS without calcium or magnesium (Gibco, #14190136) containing protease (Sigma, #S8830) and phosphatase inhibitors (Sigma, #P5726) (referred to as DPBS buffer). After final wash (see above), hepatocytes were resuspended in room temperature DPBS buffer at concentration of 1 million cells/mL and formaldehyde (ThermoFisher, #28908) was added to final concentration of 1%. Hepatocyte suspension was fixed at room temperature for 10 min with gentle rolling. Fixation was stopped by adding glycine in water (final concentration of 125 mM) containing protease and phosphatase inhibitors, and hepatocytes were incubated for additional 15 min at room temperature with gentle rolling. Hepatocytes were washed 1X in ice-cold DPBPS buffer, aliquoted in Eppendorf tubes at 1 million cells per tube, pelleted/aspirated, and snap frozen in liquid nitrogen for long-term storage at -80°C.

#### Immunoprecipitation

Immunoprecipitation reagents was prepared by washing Protein A Dynabeads (ThermoFisher, #10002D) in chromatin dilution buffer (0.5% Triton X-100, 2mM EDTA, 20mM Tris, 150 mM NaCl in water) on DynaMag-2 (ThermoFisher, #12321D) and combining with anti RNAPII subunit B1 (Rpb1) NTD rabbit monoclonal antibody (Cell Signaling, #14958) at ratio of 630 µg Dynabeads to 4 µg antibody per IP reaction. Dynabead-antibody mixture was pre-incubated at 4°C for 3 hours with gentle rolling.

#### Sonication

Fixed-frozen hepatocyte pellets (1 million cells per IP reaction) were thawed on ice and washed 1X with 500 µL cold DPBS buffer before resuspension in 500 µL cold DPBS buffer with 0.5% Triton-X100 and incubation on ice for 10 min. Hepatocytes were then centrifuged at 2600 xg for 5 min at 4°C, gently aspirated, and washed 1X with 500 µL cold tris-EDTA buffer (Sigma, #93283) containing 0.5% SDS (Sigma, #71736) with protease/phosphatase inhibitors (TE-SDS buffer) before centrifuging again at 2600 xg for 5 min at 4°C. Supernatant was carefully aspirated, pellet was resuspended with 200 µL cold TE-SDS buffer, and sample was transferred to 1.5 mL Bioruptor Pico tubes (Diagenode, #C30010016) on ice before sonication in 4°C water bath for 3 cycles (30 sec on / 30 sec off) with Easy Mode using Bioruptor Pico sonication device (Diagenode, #B01080010). After sonication, sample was transferred back to 1.5 mL Eppendorf tubes on ice and combined with 800 µL cold chromatin dilution buffer with protease/phosphatase inhibitors (see above). Samples were centrifuged at 16,000 xg for for 20 min at 4°C to remove debris and supernatant containing soluble chromatin was transferred to new Eppendorf tubes on ice. DNA concentration across all samples was checked using Qubit dsDNA HS kit (ThermoFisher, #Q33231) and normalized with chromatin dilution buffer. For each sample, a 200 µL aliquot was collected as input and a second aliquot (900 µL) was combined with 100 µL of Dynabead-antibody mixture, briefly vortexed, and incubated overnight at 4°C with gentle rolling.

#### Washing

After overnight incubation, IP samples were washed in 4°C cold-room as follows. Samples were placed on DynaMag-2 for 1 min, supernatant was removed and discarded, 500 µL of pre-chilled wash buffer was added, and samples were rotated end-over-end in cold-room for 5 min before repeating this process. Washes consisted of first RIPA-140 (0.1% SDS, 1% Triton X-100, 1 mM EDTA, 10 mM Tris, 140 mM NaCl, 0.1% sodium deoxycholate (Sigma, #D6750) in water), second RIPA-300 (0.1% SDS, 1% Triton X-100, 1 mM EDTA, 10 mM Tris, 300 mM NaCl, 0.1% sodium deoxycholate in water), third LiCl buffer (250 mM LiCl (Fisher Scientific, #L121), 0.5% Igepal ca-630 (MP Biomedicals, # 198596), 1 mM EDTA, 10 mM Tris, 0.5% sodium deoxycholate in water), fourth Tris-EDTA buffer (Sigma, #93283) with 0.2% Triton X-100, and fifth Tris-EDTA buffer alone.

#### Elution

Wash buffer was removed, and chromatin was eluted from Dynabeads as follows^37^. Briefly, 100 µL of elution buffer (1% SDS, 1 mM EDTA, 10 mM Tris in water) was added to beads before vortexing and incubating at 65°C for 5 min followed by room temperature incubation for 15min with vortexing every 5 min. Beads were placed on DynaMag-2 for 1 min, and supernatant containing immunoprecipitated chromatin (∼100 µL) was carefully collected and saved. Beads then underwent a second round of elution, and the two supernatant collections were combined (∼200 µL total).

#### RNase digestion, reverse cross-linking, and DNA purification

Input and IP samples were combined with NaCl (final concentration of 160 mM) and RNase A (final concentration of 20 µG/mL) (Roche, # 11119915001) and incubated at 65°C overnight as previously outlined^37^. Afterwards, EDTA (final concentration of 5 mM) and proteinase K (final concentration 200 µG/mL) (Invitrogen, # 100005393) were added, and samples were incubated at 45°C for 2 h. Finally, input and IP DNA was purified using ChIP DNA Clean & Concentrator kit (Zymo Research, #D5205) according to manufacturer’s guidelines, and DNA concentration was quantified using Qubit dsDNA HS kit. Fraction of purified input DNA was run on agarose gel to assess fragmentation pattern (∼100-500 bp).

#### Library preparation, sequencing, and data processing

Library preparation was performed using NEBNext® Ultra™ II DNA Library Prep Kit for Illumina® (New England Biolabs, #E7645) according to manufacturer’s guidelines with NEBNext® Multiplex Oligos for Illumina® single Index Primers Set 1 and Set 2 (New England Biolabs, #E7335 and #E7500). Premade libraries were sent to Novogene for sequencing using NovaSeq paired-end sequencing at 150 base pairs per read (PE150). Raw fastq files were first processed to remove adapter sequences and low-quality bases using Trim Galore (v0.6.7) with “--stringency 5”. Trimmed reads were then aligned to mm10 reference genome using bowtie2 (v2.4.5) with default parameters^34^. The resulting SAM files were converted to BAM files using samtools (v1.6) view with “-f 2 -q 10”^35^. Duplicates were removed using samtools rmdup. Coverage tracks were generated using the bamCoverage tool from the deepTools (v3.5.1) suite with the “--extendReads - -normalizeUsing CPM” option^36^. Input normalization was performed using log2 ratio and bin size set to 5 with bigwigCompare from deepTools. Normalized BigWig files were visualized using Integrative Genomics Viewer (IGV)^38^. Total hepatocyte gene tracks (Fig. 1 and Fig. S1) represent IP over input signal; periportal/pericentral hepatocyte gene tracks (Fig. S2) represent IP signal alone since input signal was negligible in previous experiments.

### RNA isolation and quantitative PCR (RT-qPCR)

Total RNA isolation from isolated hepatocytes was done using the RNeasy mini kit (Qiagen, #74106) in combination with TRIzol (Invitrogen, #15596026) according to the manufacture’s protocol. cDNA was synthesized using SuperScript IV VILO (Invitrogen, #11756050) and RT-qPCR was performed using PowerUp SYBR Green Master Mix (Applied Biosystems, #A25778) on a QuantStudio 6 system (Applied Biosystems). Relative gene expression was determined by the ΔΔCt method and normalized to *Ppib*. Primer sequences are listed in Table S2.

### Targeted Single-cell RNA sequencing (targeted scRNA-seq)

To characterize dual-modal hepatocytes, we analyzed a publicly available targeted single-cell RNA sequencing (scRNA-seq) dataset covering 61 specific genes^5^. The panel consisted of pericentral and periportal hepatocyte markers, general parenchymal and non-parenchymal cell markers, as well as genes involved in gluconeogenesis, lipogenesis, bile acid synthesis, and several secreted hepatokines. Raw FASTQ files were obtained from the NCBI Gene Expression Omnibus (GEO; accession no. GSE263419). Gene expression levels were quantified using Cell Ranger (v.6.0.1; 10x Genomics) in targeted gene expression mode. Positive cells were defined by applying thresholds determined via ridge plot analysis and compared with RT-qPCR results as previously described^5^.

### HCR RNA-FISH

Livers were fixed in 4% paraformaldehyde (PFA) for 3 hours, cryoprotected in PFA + 30 % sucrose for overnight, embedded in OCT, and tissue blocks were kept at -80 °C until sectioning. After sectioned at 8-10 μm thickness, the sections were post-fixed, permeabilized with ethanol, and treated with proteinase K prior to hybridization as previously described^5^. HCR RNA-FISH probes (Molecular Instruments) targeting *Pck1* (B3), *Fasn* (B1), and *Glul* (B2) were hybridized overnight at 37 °C (0.4 pmol of each probe set/100 uL). Once excess probes were removed, the amplification stage was performed according to the manufacture’s protocol for 12 hours (B3=546, B1=647, B2=488, 6 pmol each of hairpin/100 uL)^39^. Sections were then washed, counterstained with DAPI, and mounted with antifade medium. Images were acquired on an upright, wide-field Olympus BX- 63 Microscope equipped with a Super Apochromatic objective lens (60×/1.35 N.A., Pixel size (x,y) = 107.5 nm, z-step = 300nm). MetaMorph software (Molecular Devices, v7.10.1.161) was used for controlling microscope automation and image acquisition. Pericentral hepatocytes were identified by *Glul* expression, and periportal hepatocytes were identified by the bile duct-associated morphology. Liver lobule images were taken by taking sequential images and processed on FIJI (v2.16.0). Transcription site analysis in hepatic lobules was performed using SmallFish (v2.1.10), a python implementation of FishQuant^40^. Cell segmentation was performed in 2D using Cellpose (https://github.com/MouseLand/cellpose); and spot detection was performed via big-fish package (https://github.com/fish-quant/big-fish).

### Stable isotope tracing analyses

For pyruvate-to glucose tracing experiments, isolated hepatocytes (1×10^5^ cells) were incubated in KH buffer containing 1 mM [2,3-^13^C_2_] sodium pyruvate or unlabeled pyruvate, 1 mM sodium lactate, 2 mM L-glutamine, and 1 mM BSA-conjugated palmitic acid at 37 °C for 1 hour. 200 μL of the media was extracted with methanol containing 2-deoxyglucose, centrifuged (13,100 × *g*, 10 min, 4 °C), and the supernatant was dried under gentle nitrogen flow. Samples were then reacted with hydroxylanmine followed by acetic anhydride as previously described^5^. After reconstitution in ethyl acetate, the samples were analyzed by gas chromatography mass spectrometry (GC (7890)-MS (5975C), Agilent) in chemical ionization (CI) mode using methane as the CI gas. Mass isotopologue distributions were analyzed using MassHunter software (Agilent) and corrected for natural isotope abundance. Glucose isotopologues are reported as molar fractions (M0–M6). The sum of all isotopologues of the molecules, m_i_ (where i = 1 to 6), is equal to 1 or 100%. The labelled isotopologue fractions consist of a distribution not affected by the dilution with unlabeled compounds, reported as m_i_/Σm_i_, in percent. The enrichment ΣM_n_ is the weighted sum of the labelled species (ΣMn = 1*M_1_ + 2*M_2_ + 3*M_3_ + 4*M_4_ + 5*M_5_ + 6*M_6_) = average number of ^13^C carbons/molecule as previously reported^5^.

For *in vivo* deuterium labeling, mice were subjected to time-restricted feeding as described above. For fed conditions, mice received an intraperitoneal (IP) injection of 0.9% NaCl in deuterated water (30 μL/g body weight) at 07:00, after which drinking water was replaced with 6% ^2^H_2_O. Chow and 6 % ^2^H_2_O with 20 % sucrose (w/v) were provided at 09:00, and mice were sacrificed at 13:00. For fasted conditions, mice were fed chow and sucrose water (without ^2^H_2_O) from 09:00 to 13:00, fasted for 24 h, injected with ^2^H_2_O and regular drinking water was replaced with 6 % ^2^H_2_O water at 07:00 the following day, and sacrificed at 13:00. Lipids were extracted from isolated hepatocytes (1×10^6^ cells) following alkaline saponification in 1 M NaOH in 70 % ethanol in water with heptadecanoic acid (C17:0) as an internal standard. Extracted fatty acids were derivatized with BSTFA and analyzed by GC-MS (Agilent) with a DB-5MS column. Palmitate deuterium enrichment was quantified from the m/z 313 fragment and calculated. The total labeled fraction of palmitate was calculated with fraction enrichment. The total palmitate was calculated relative to the concentration of heptadecanoic acid. The newly synthesized palmitate was calculated with the function: Newly synthesized palmitate = Total palmitate * total fraction of labeled palmitate/deuterium enrichment in plasma/tissue weight.

### 18O water labeling in vivo

Mice were subjected to time-restricted feeding as described above. In both fed and fasted conditions, the mice were gavage with ^18^O water or regular water at 12:48, received IP injection of ^18^O water or regular water at 12:53, then sacrificed at 13:00. The liver was rinsed with 150 mM ammonium acetate before frozen in pre-chilled isopentane. A 20 um thickness of tissue section was used for DESI imaging. The slide was analyzed with a Waters Cyclic Ion Mobility-Mass Spectrometry (Cyclic-iMS). The slide was placed in desiccator for about half an hour before loading into DESI stage. The solvent was methanol:water=60:40(v:v) with 200ng/mL LeuEnk. The flow rate was set to 2 µL/min. The resolution was set to 60um with 300 µm/sec. The mass range was set to 50 – 1200 m/z, and the mobility was set to 1 pass with TW ramp of TOF settings are 16 – 28 V/ms.

### DESI and MALDI metabolic spatial imaging

Mice were subjected to time-restricted feeding as described above. On the day of the experiment, mice were provided with D17011901px (Modified OSD with Macronutrient Content Similar to LabDiet 5053) with ^13^C_6_ glucose or unlabeled glucose (for 1 g of premix, 1.31 g of glucose was added) at 9:00. Mice were then sacrificed at 13:00, corresponding to 4 hours of feeding, and the liver was frozen in pre-chilled isopentane. For DESI analysis, the slices were sectioned into 20um thickness. The ^13^C samples was analyzed with DESI-Xevo TQ absolute MS (Waters). The solvent was methanol:water=80:20(v:v) with 200ng/mL LeuEnk with a flow rate of 1.2ul/mL and was scanned with 25 x 25 um spatial resolution. The metabolites and their isotopolgues was analyzed in the negative multiple reaction monitoring (MRM) mode. The data was processed with HDI software (Waters).

### PrimeFlow

After isolating hepatocytes as described above, the hepatocytes went through the PrimeFlow (Invitrogen, #88-18005-204) protocol based on the manufacturer’s instructions with minor modifications. Briefly, hepatocytes were fixed using PrimeFlow RNA Fixation Buffers. Cells were then permeabilized in PrimeFlow RNA Permeabilization Buffer supplemented with RNase inhibitors, followed by a second fixation step. After washing, cells were hybridized with target-specific RNA probe sets diluted in PrimeFlow RNA Target Probe Diluent for 2 hours at 40 °C. Samples were washed in PrimeFlow RNA Wash Buffer containing RNase inhibitor and stored overnight at 2-8 °C in the dark. On the following day, signal amplification was performed through sequential incubations with PreAmp Mix, Amp Mix, and fluorescently labeled probes, each at 40 °C, with intervening washes according to the manufacturer’s instructions. Following the final labeling step, cells were washed, resuspended in PrimeFlow RNA Storage Buffer and analyzed on BD FACSAria III. All centrifugation steps were done at 800 × *g* unless otherwise specified.

### Hyperinsulinemic-euglycemic clamp experiments

Hyperinsulinemic-euglycemic clamp studies were performed in conscious, unrestrained mice. Briefly, mice were surgically implanted with catheters in the jugular vein for infusions and the carotid artery for blood sampling. After recovering for at least 3 days following surgery, the mice were subjected to time-restricted feeding as described above. On the day of the experiment, catheters were connected to infusion lines, and basal blood glucose was measured from the arterial blood using a glucometer. Continuous insulin was infused at the indicated rates and maintained for 2 hours. Blood glucose was clamped at 150 mg/dL by a variable infusion of 20% glucose solution, adjusted based on blood glucose measurements every 10-20 minutes. At the end of the clamp, liver perfusion for total hepatocyte isolation was performed.

### Spatial Transcriptomics

For high-fat diet studies, fresh frozen mouse livers were embedded in Neg-50 and stored at -80 °C until sectioning. After sectioned at 10 μm thickness, **s**ections were adhered to capture areas by brief warming onto Visium v1 slides (10x Genomics). For humanized liver studies, livers were fixed in 4% paraformaldehyde (PFA) for 3 hours, cryoprotected in PFA + 30 % sucrose for overnight, embedded in OCT, and tissue blocks were kept at -80 °C until sectioning. After sectioned at 10 μm thickness, Visium v2 (prepared with CytAssist) was performed using the CytAssist instrument. For both studies, the sample preparation was performed according to the manufacturer’s guidelines. Briefly, tissue sections were stained with hematoxylin and eosin (H&E), imaged using a brightfield microscope for histological reference and spatial registration. RNA capture, reverse transcription, second-strand synthesis, cDNA amplification, and library construction were performed. Libraries were indexed, quality-controlled, and sequenced on an Illumina platform using paired-end sequencing. Raw sequencing data were processed using Space Ranger (v2.0.0 for high-fat diet studies with reference genome: mm10-2020-A, v3.1.0 for humanized liver studies with reference genome: GRCh38-2020-A, 10x Genomics) to align reads to the reference genome and generate spatial gene expression matrices. Spatial alignment was performed using H&E images, and downstream analyses were conducted using standard single-cell and spatial transcriptomics workflows.

## QUANTIFICATION AND STATISTICAL ANALYSIS

Excel (v16.101.3), Prism (v10.6.1) GraphPad Software, were used for data processing, analyses, and graph productions in the experiments. All data are presented as the mean +/- standard deviation unless otherwise noted. Student’s *t*-test with unpaired two-tailed p values of *p<0.05, **p<0.01, and ***p<0.001 was considered statistically significant when comparing two groups. (N.S.=not statistically significant). Figures/Images were created by Powerpoint (v16.101.3), IGV (v2.19.7), Loupe Browser (v9.0.0), FIJI (v2.16.0), QuPath (v0.6.0) and Adobe Illustrator (v28.1).

## Supplementary Figures

**Figure S1.**
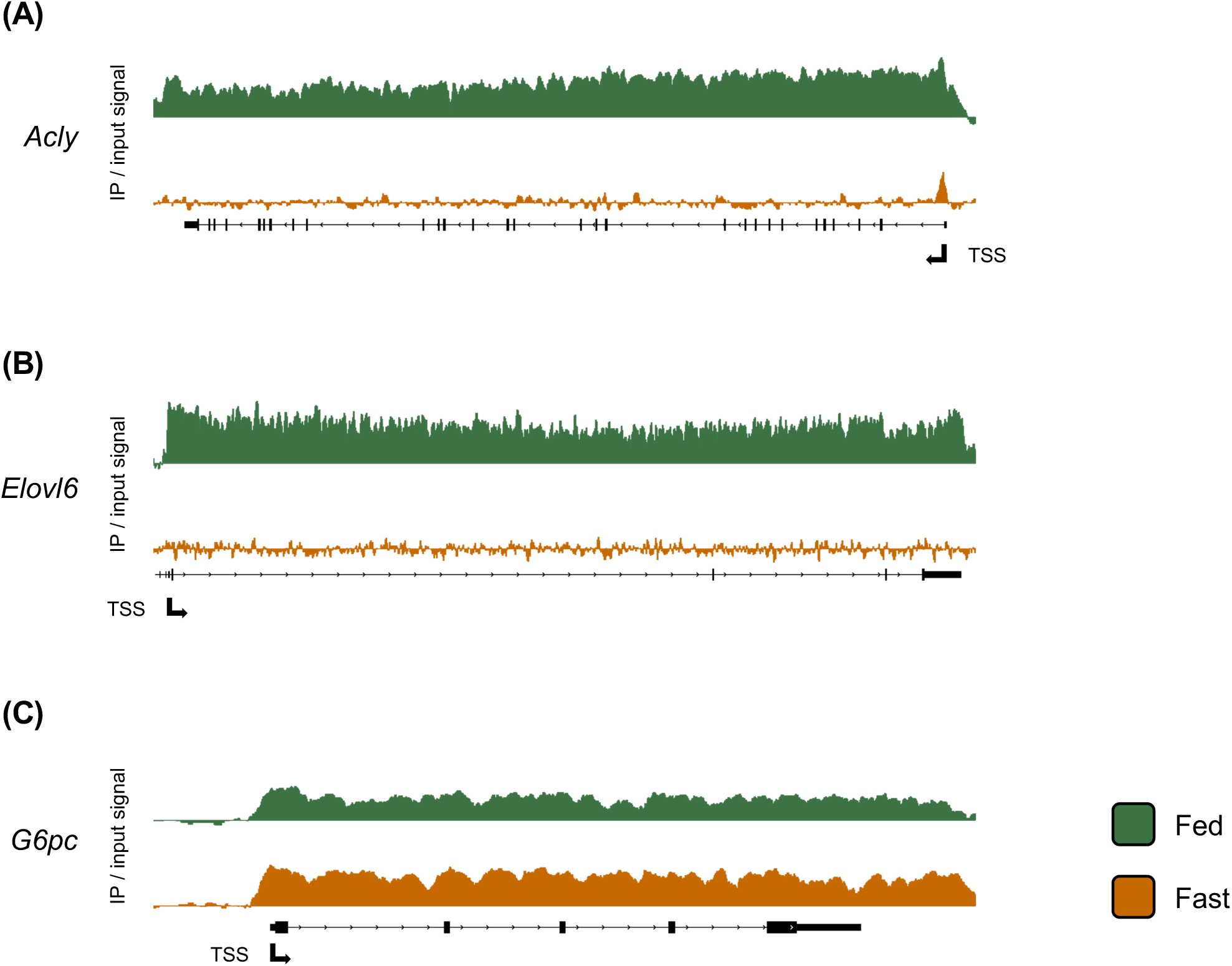
*G6pc* is transcribed in hepatocytes in the fed state. (A) RNAP II ChIP of *Acly* in fed/fasted total hepatocytes. (B) RNAP II ChIP of *Elovl6* in fed/fasted total hepatocytes. (C) RNAP II ChIP of *G6pc* in fed/fasted total hepatocytes.

**Figure S2.**
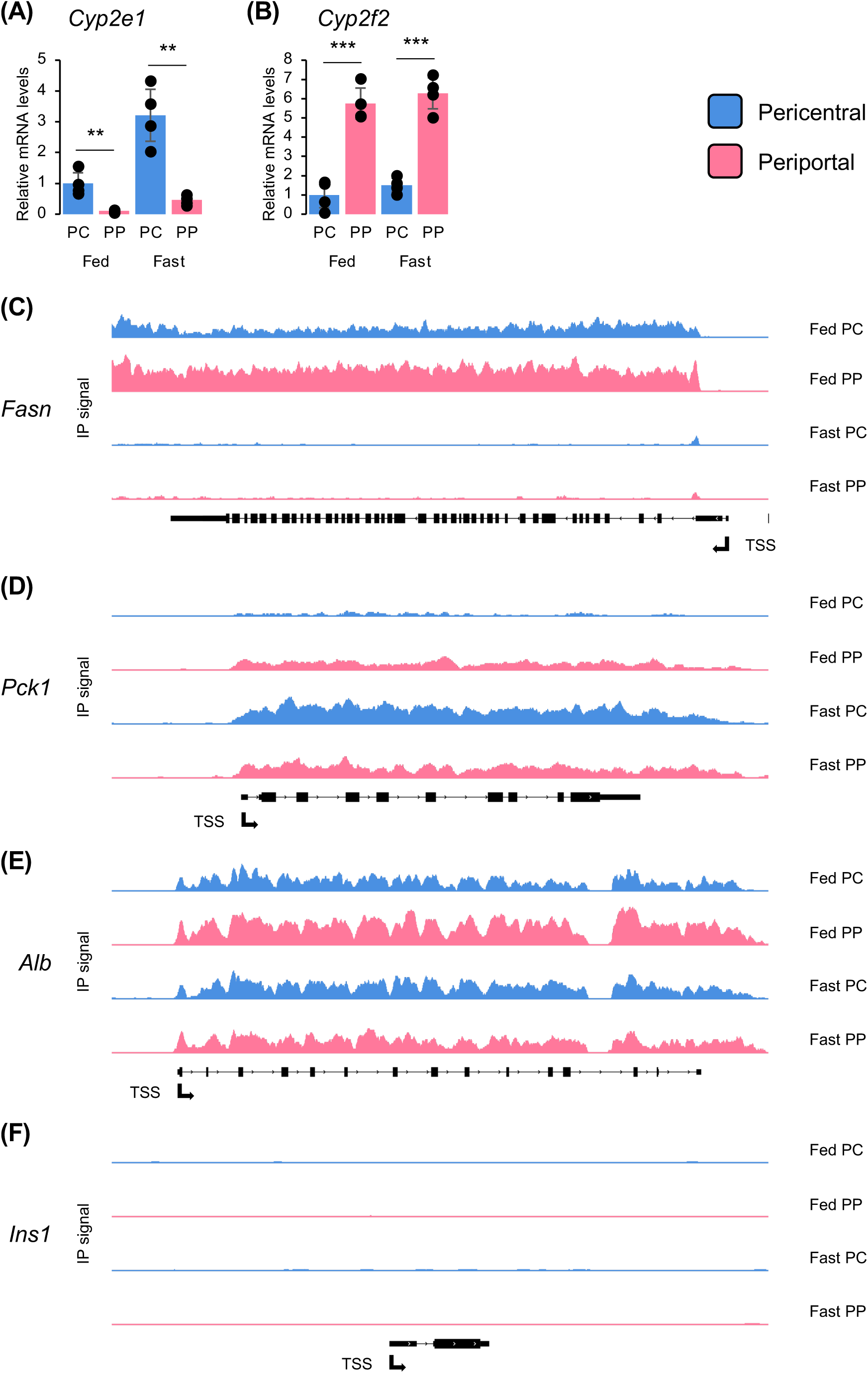
*Pck1* is transcribed in periportal hepatocytes in the fed state. (A) Relative mRNA levels of *Cyp2e1* in in fed/fasted PC/PP hepatocytes. (B) Relative mRNA levels of *Cyp2f2* in in fed/fasted PC/PP hepatocytes. RNAP II ChIP of *Fasn* in fed/fasted PC/PP hepatocytes. (D) RNAP II ChIP of *Pck1* in fed/fasted PC/PP hepatocytes. (E) RNAP II ChIP of *Alb* in fed/fasted PC/PP hepatocytes. (F) RNAP II ChIP of *Ins1* in fed/fasted PC/PP hepatocytes.

**Figure S3.**
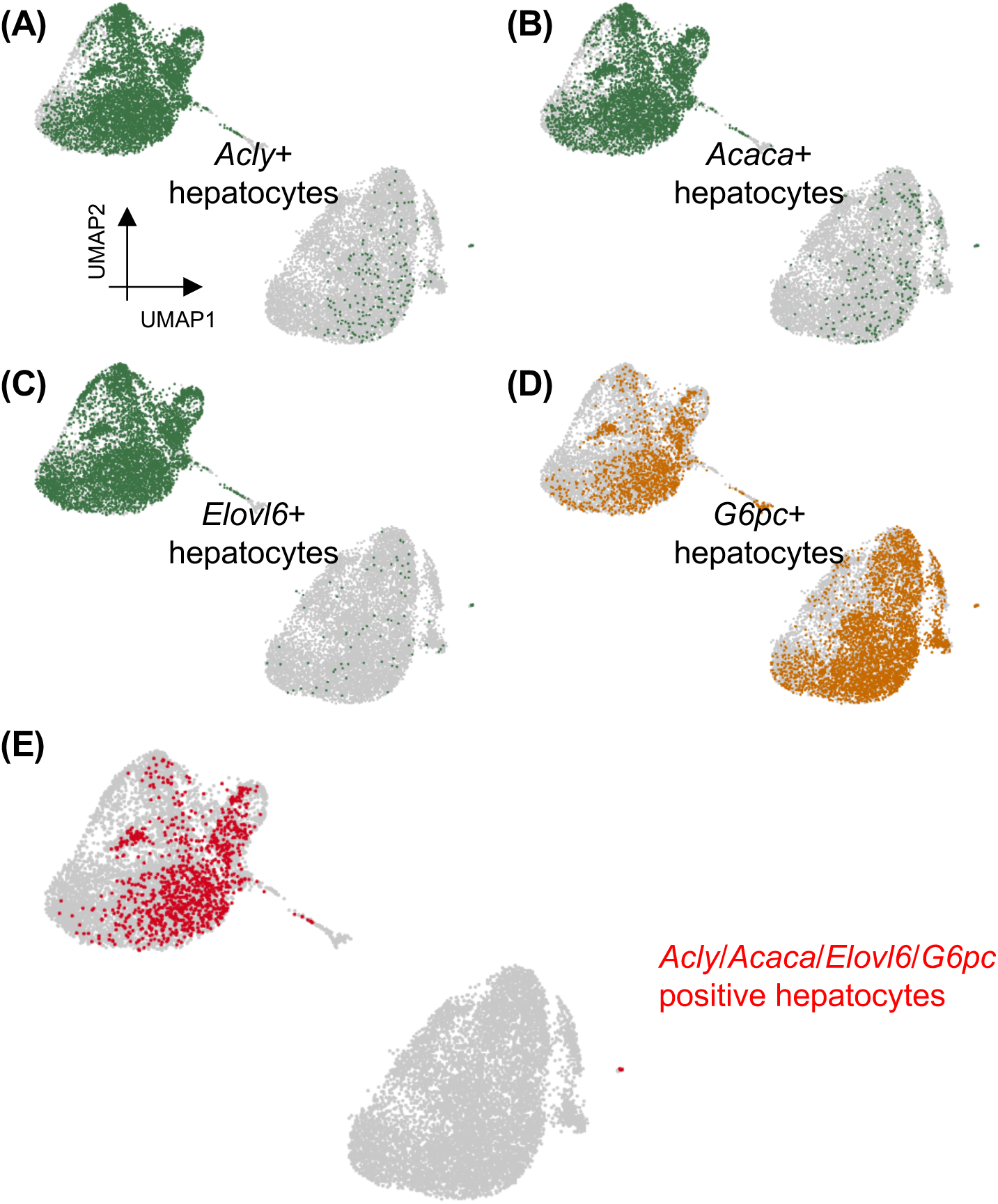
*Acly*, *Acaca*, *Elovl6*, and *G6pc* are co-expressed in a subset of hepatocytes. (A) UMAP of *Acly* positive hepatocytes. (B) UMAP of *Acaca* positive hepatocytes. (C) UMAP of *Elovl6* positive hepatocytes. (D) UMAP of *G6pc* positive hepatocytes.

**Figure S4.**
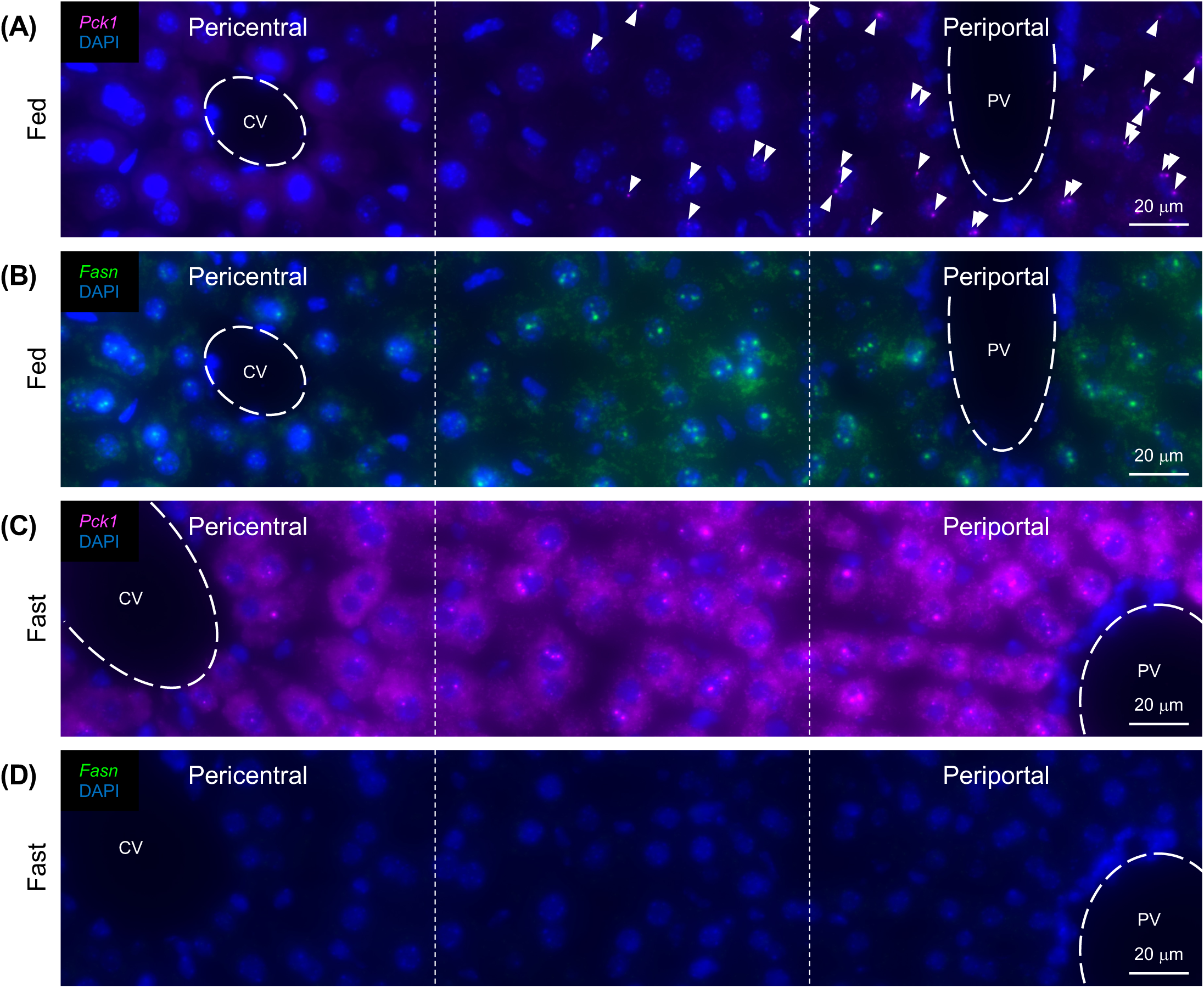
*Pck1* is actively transcribed in the fed state. (A) HCR RNA-FISH image of *Pck1* (magenta) with DAPI-stained nuclei (blue) in the fed state (representative image shown). Lobules show the central vein (CV) on the left and the portal vein (PV) on the right. (B) HCR RNA-FISH image *Fasn* (green) with DAPI-stained nuclei (blue) in the fed state (representative image shown). Lobules show the central vein (CV) on the left and the portal vein (PV) on the right. (C) HCR RNA-FISH image of *Pck1* (magenta) with DAPI-stained (blue) in the fasted state (representative image shown). Lobules show the central vein (CV) on the left and the portal vein (PV) on the right. (B) HCR RNA-FISH image *Fasn* (green) with DAPI-stained (blue) in the fed state (representative image shown). Lobules show the central vein (CV) on the left and the portal vein (PV) on the right. Scale bar, 20 um.

**Figure S5.**
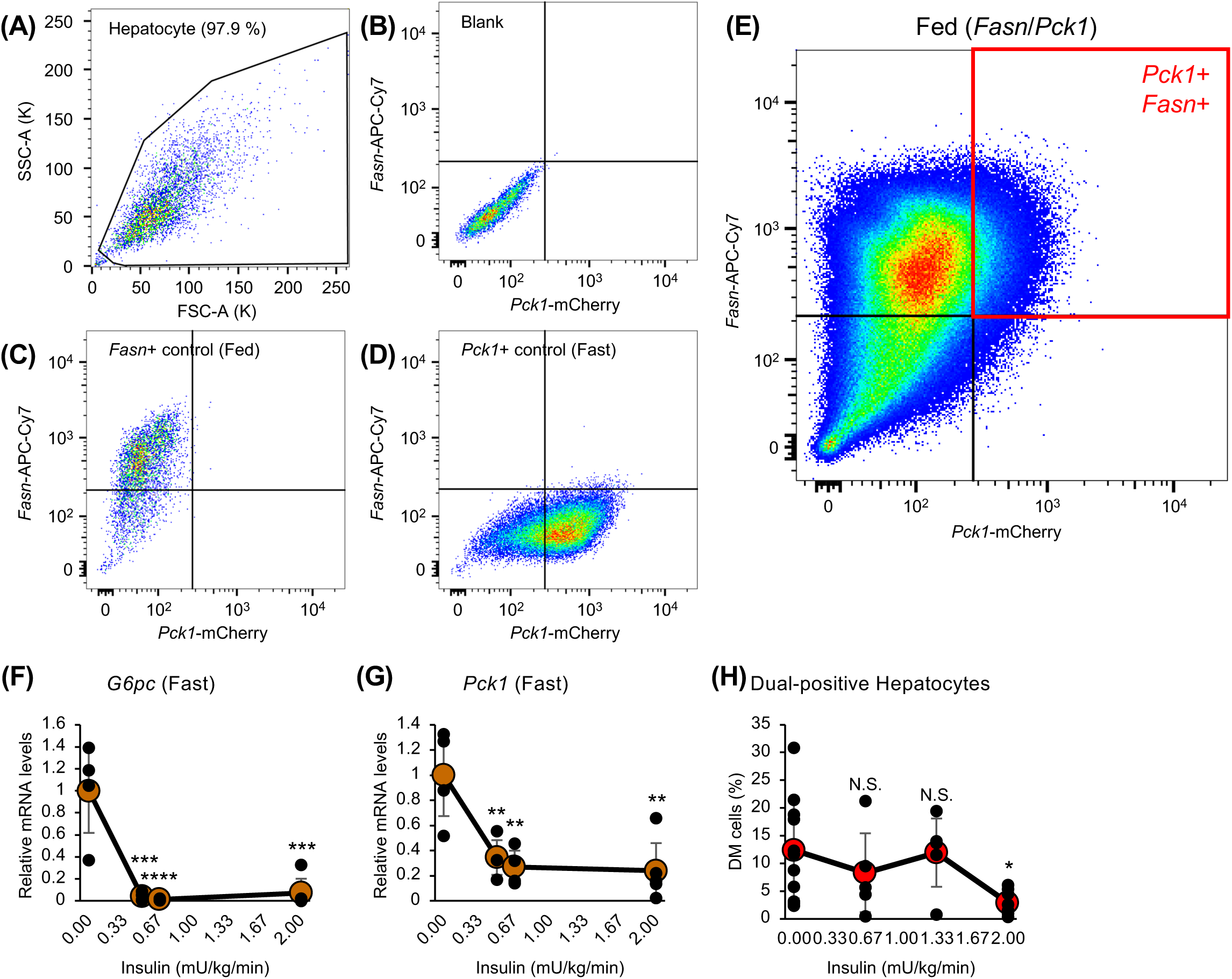
Dual-positive hepatocytes are insulin resistant in suppression of *Pck1*. (A) Flow cytometry gating strategy utilizing FSC-A and SSC-A for size selection of hepatocytes. (B) Background fluorescence of hepatocytes. (C) Gating of *Fasn* based on fed hepatocytes. (D) Gating of *Pck1* based on fasted hepatocytes. (E) Representative flow cytometry result of *Fasn*/*Pck1* in the fed state; the red quadrant shows the Dual-modal (DM) hepatocytes. (F) Insulin suppression of *G6pc* mRNA expression based on hyperinsulinemic-euglycemic clamps on fasted hepatocytes. (G) Insulin suppression of *Pck1* mRNA expression based on hyperinsulinemic-euglycemic clamps on fasted hepatocytes. (H) Insulin suppression of *Pck1* mRNA expression on DM hepatocytes based on hyperinsulinemic-euglycemic clamps in the fed state.

**Figure S6.**
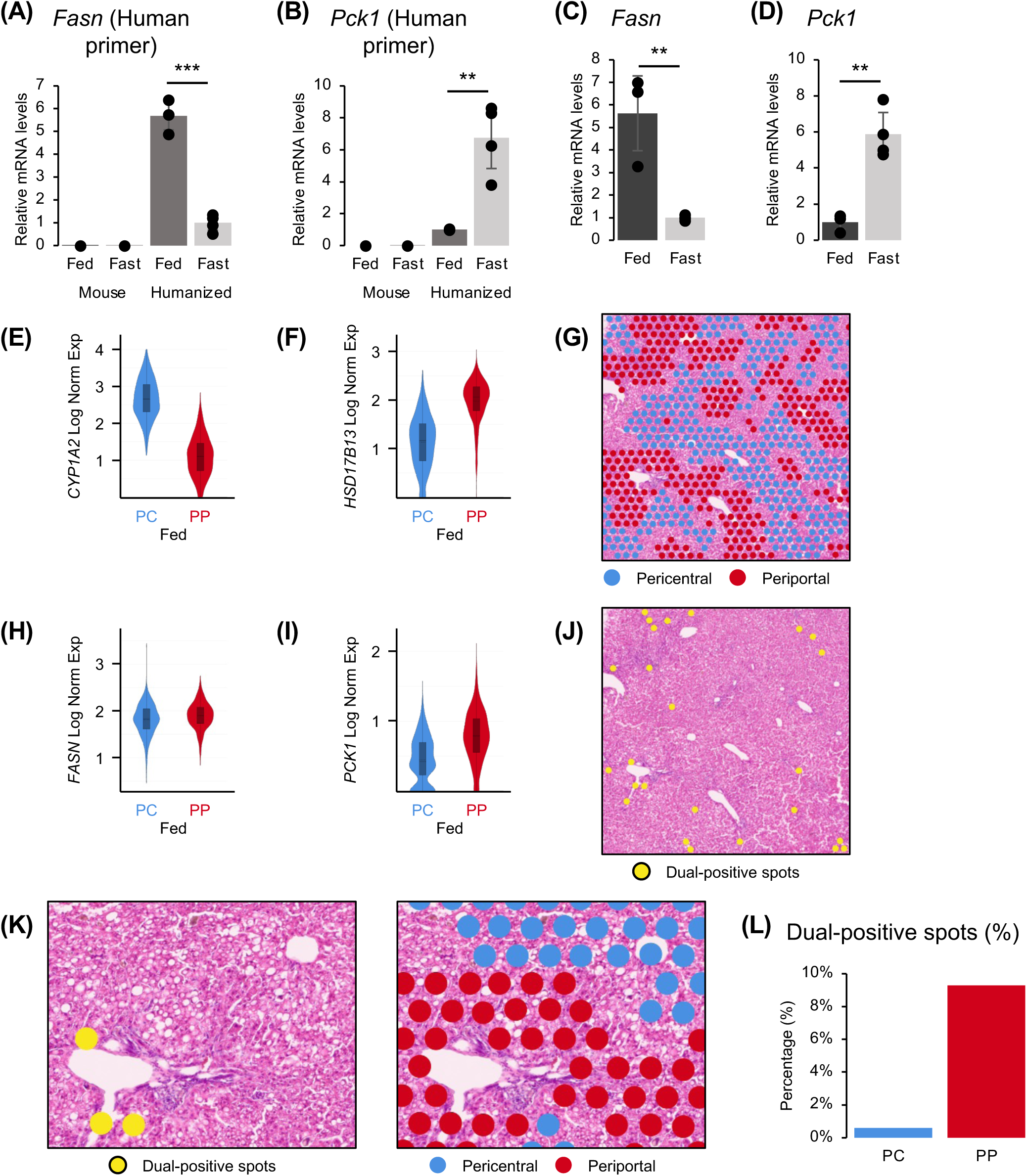
Humanized mouse liver has dual-positive spots. (A) Relative mRNA levels of human *Fasn* from fresh frozen liver tissue (mouse liver vs. humanized liver, fed/fasted). (B) Relative mRNA levels of human *Pck1* from fresh frozen liver tissue (mouse liver vs. humanized liver, fed/fasted). (C) Relative mRNA levels of human *Fasn* from 4% PFA fixed liver sections (humanized liver). (D) Relative mRNA levels of human *Pck1* from 4% PFA fixed liver sections (humanized liver). (E) Violin plot of *Cyp1a2* (PC marker) log normalized expression. (F) Violin plot of *Hsd17b13* (PP marker) log normalized expression. (G) Spatial Transcriptomics slide from fed mice with Pericentral (PC)/Periportal (PP) spots highlighted. (H) Violin plot of *Fasn* (lipogenic gene) log normalized expression. (I) Violin plot of *Pck1* (gluconeogenic gene) log normalized expression. (J) Spatial Transcriptomics slide from fed mice with Dual-Modal (DM) spots highlighted. (K) DM spots image with PC/PP alignment. (L) Percentage of DM spots in PC/PP regions.

**Figure S7.**
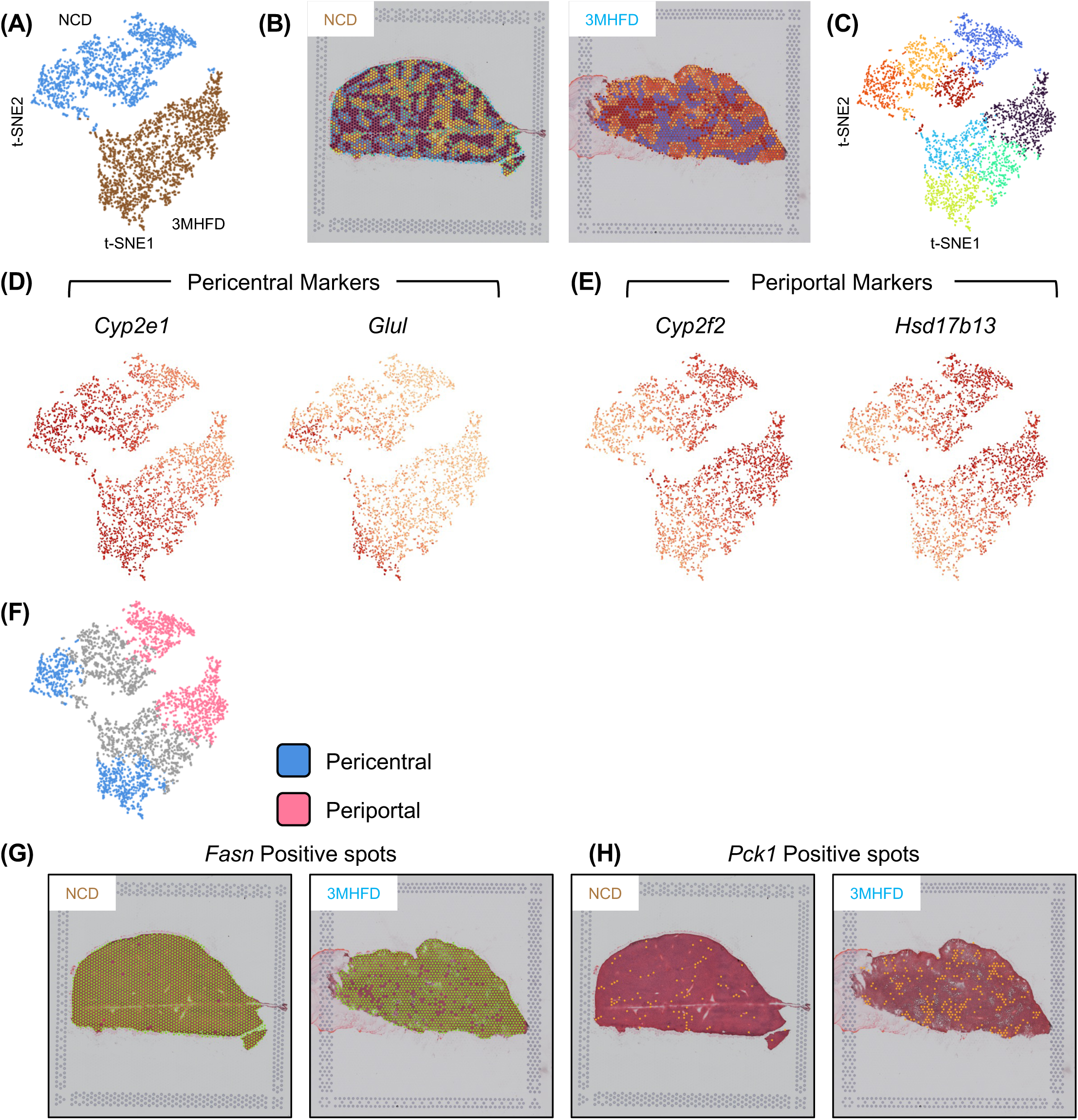
t-SNE separates NCD and 3MHFD. (A) t-SNE visualization of spatial transcriptomics based on feeding conditions (Normal Chow Diet (NCD) and 3-Month High-Fat Diet (3MHFD). (B) Spatial transcriptomics image of NCD fed mice and 3MHFD fed mice with graph-based clustering highlighted. (C) t-SNE visualization of spatial transcriptomics based on graph-based clustering. (D) t-SNE visualization of pericentral markers *Cyp2e1* and *Glul*. (E) t-SNE visualization of periportal markers *Cyp2f2* and *Hsd17b13*. (F) t-SNE visualization of spatial transcriptomics based on zonation. (G) spatial transcriptomics image of *Fasn* positive spots in NCD and 3MHFD fed mice. (H) spatial transcriptomics image of *Pck1* positive spots in NCD and 3MHFD fed mice.

**Table S1.**
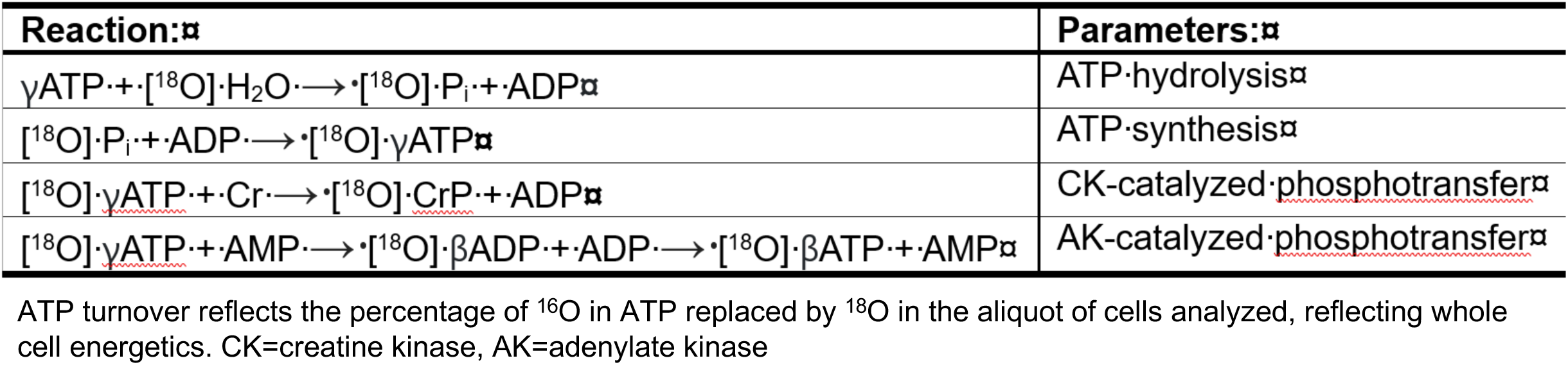
Compendium of ^18^O/^16^O exchange reactions which determine ATP turnover.

**Table S2.**
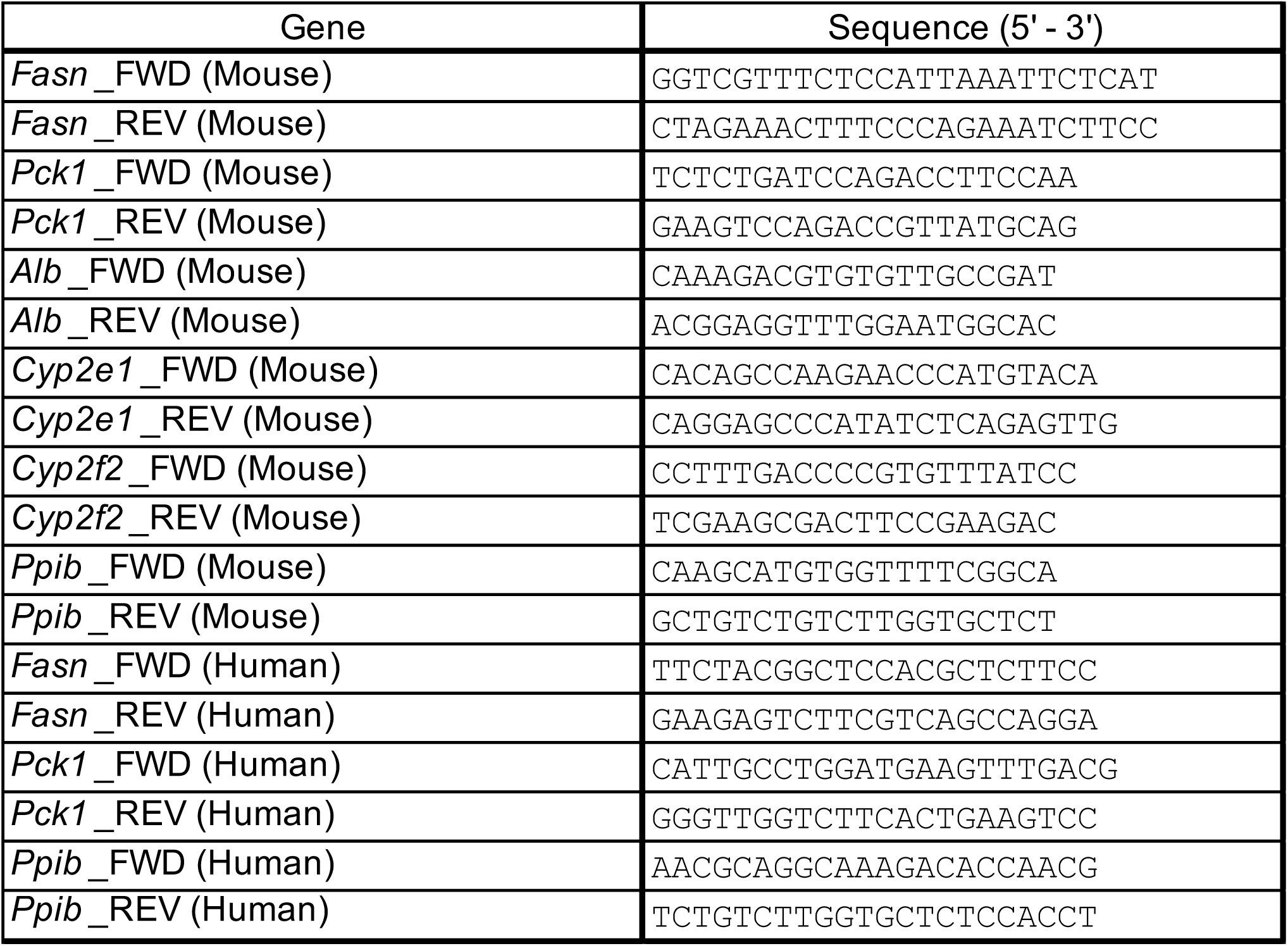
Primer Sequences.

